# The forgotten psychedelic: Spatiotemporal mapping of brain organisation following the administration of 2C-B and psilocybin

**DOI:** 10.1101/2024.10.22.619393

**Authors:** Pablo Mallaroni, Parker Singleton, Natasha L. Mason, Theodore D. Satterthwaite, Johannes G. Ramaekers

**Author notes:** **Corresponding authors’ emails**.

## Abstract

As psychedelic-assisted psychotherapy gains momentum, clinical investigation of next-generation psychedelics may lead to novel compounds tailored for specific populations. 2,5-dimethoxy-4-bromophenethylamine (2C-B) is a psychedelic phenethylamine reported to produce less dysphoria and subjective impairment than the psychedelic tryptamine psilocybin. Despite its popularity among recreational users and distinct pharmacodynamics, the neural correlates of 2C-B remain unexplored. Using 7T resting-state functional MRI in 22 healthy volunteers, we mapped out the acute effects of matched doses of 20 mg 2C-B, 15 mg psilocybin and placebo across spatiotemporal benchmarks of functional brain organisation. In a within-subjects, double-blind, placebo-controlled crossover design, we evaluated the neuropharmacological and neurobehavioural correlates of an array of connectivity measures – including static (sFC) and global connectivity (gFC), dynamic connectivity variability (dFC), and spontaneous brain complexity. Compared to placebo, 2C-B and psilocybin selectively reduced intra-network sFC, while broadly increasing between-network and subcortical-cortical connectivity. Compared to psilocybin, 2C-B exhibited less pronounced reductions in between-network FC but elicited elevations in transmodal sFC. Both compounds yielded spatially divergent increases in gFC yet produced similar increases in brain complexity. Using PET density modelling, the spatial distribution of neural effects aligned with documented differences in monoaminergic transporter and serotonergic receptor binding affinity beyond 5-HT_2A_, highlighting the role of pharmacology in shaping functional dynamics. Lastly, we show behavioural markers of psychedelic effects are non-linearly reflected by the desynchronisation of the transmodal axis of functional brain organisation. Together, our findings highlight 2C-B as a useful new addition to the study of psychedelic neuroscience and may motivate new pharmacotherapy strategies.

## Introduction

There is currently intense clinical interest in the use of psychedelics for a range of neuropsychiatric disorders. Current research suggests that “classical” candidate compounds such as psilocybin partly exert their therapeutic effects by temporarily disrupting maladaptive functional brain organisation via 5-HT_2A_ agonism (1, 2). Several human resting-state functional magnetic resonance imaging (rsfMRI) studies have reported classical psychedelics acutely decrease blood-oxygen-level-dependent (BOLD) signal variance and broadly increase functional connectivity (FC) across the neocortex (3, 4). These findings underlie several theories implicating reduced functional network segregation and increased neural complexity (often referred to as entropy) as the basis of psychedelic action (5). As the field advances, there is now growing attention towards next-generation psychedelics which may elicit similar cortical effects (6). Developing compounds that induce milder, shorter, or more tolerable altered states of consciousness may be more feasible to scale in clinical practice while offering new mechanistic insights due to diverging pharmacodynamics (7, 8).

One such compound is the novel phenethylamine 4-bromo-2,5-dimethoxyphenethylamine (2C-B). While 2C-B is the most frequently used novel psychedelic among recreational drug users (9) and has a history of therapeutic use (10), scarce clinical research data exists on its effects. Compared to the prototypical psychedelic tryptamine psilocybin, 2C-B appears to induce less subjective impairment and dysphoria at comparable doses, while producing similar perceptual and cognitive effects (11, 12), making it potentially easier to tolerate. A partial agonist, 2C-B engages multiple serotonin receptors beyond 5-HT_2A_, including 5-HT_1A_, 5-HT_1B_ and 5-HT_2C_ while sharing a similar primary binding profile to psilocin, the active biometabolite of psilocybin (13–16). Experiential differences between these two compounds may be attributed to their functional selectivity for 5-HT_2A_. receptors; with 2C-B showing approximately tenfold greater selectivity for 5-HT_2A_ over 5-HT_1A_ compared to psilocybin, which may bias downstream signalling cascades (14, 15). In parallel, it is plausible that any of 2C-B’s distinct entactogenic and cortical effects are due to its reversal of monoamine transporters otherwise trivial to classical psychedelics (14, 17–19).

Polypharmacology is thus likely to contribute to distinct neural circuits being modulated by distinct psychedelics, given the differing role of monoaminergic systems on cortical signalling (20). Neuromodulatory systems tune the neural ‘gain’ of receptive neuronal populations (21) thereby modulating inter-areal communication, with pronounced effects at a network level (22) (23). Under psychedelics, changes in neural dynamics and phenomenology have been attributed to distinct serotonergic receptors (24–27). Some studies have retrospectively investigated such dependencies for brain organisation and subjective experience across psychedelics (28, 29). However, within-subject comparative studies, which minimise interstudy and intersubject variability, are still lacking. Moreover, heterogeneous rsfMRI approaches lacking independent replication have been assessed as experiential markers with varying degrees of success (30, 31). Understanding that no single measure of psychedelic brain effects is yet superior, a perhaps more cogent aim is to define focal neural regions across psychedelics that are differentially sensitive to drug-induced perturbations - regardless of the assessment method.

Here, we examined 2C-B’s effects on functional brain organisation using a double-blind, crossover design against placebo and matched doses of psilocybin as an active drug control condition in healthy volunteers using 7 Tesla rsfMRI. A first direct comparison of two structurally distinct psychedelics, we hypothesised that 2C-B would elicit alterations to spatiotemporal benchmarks of brain organisation consistent with a primary 5-HT_2A_ mechanism of action, that is – the presence of a small-world network organisation and increased neural complexity. Additionally, we hypothesised that monoamine reuptake inhibition would significantly explain topographical differences between 2C-B and psilocybin. To do so, we employed a pharmacology-guided approach leveraging positron emission tomography (PET) mapping. Finally, we investigated potential relationships between multivariate markers of brain organisation and subjective experience.

## Materials and methods

All aspects of this study (trial register NL8813) were approved (NL73539.068.20) by the Academic Hospital and University’s Medical Ethics Committee of Maastricht University. All procedures were conducted according to the Declaration of Helsinki (1964), amended in Fortaleza (Brazil, October 2013) and were in accordance with the Medical Research Involving Human Subjects Act (WMO). A permit for obtaining, storing, and administering 2C-B and psilocybin were obtained from the Dutch Drug Enforcement Administration. All participants were fully informed of all procedures, possible adverse reactions, legal rights, and responsibilities, expected benefits, and their right to voluntary termination without consequences.

### Participants

Twenty-two healthy participants (11 female) aged 19-35 years (meanl1±l1SD: 25 ± 4 years) were recruited by word of mouth and advertisement shared via the Maastricht University social media platform. Participants were required to have had at least one previous lifetime psychedelic exposure (e.g., psilocybin, mescaline, ayahuasca, LSD), but no exposure within the past 3 months. Those with psychiatric, major medical, endocrine, or neurological conditions, pregnant or lactating women, those not using reliable contraception, conconimitatant drug use, with a history of adverse reactions to psychedelics, with uncorrected or abnormal vision, or MRI contraindications were excluded (see Supplementary Materials for a full inclusion criterion and testing procedures).

### Study design

Healthy adults were enrolled in a double-blind, placebo-controlled, randomised crossover design with three acute fMRI imaging sessions to investigate the effects of 2C-B and psilocybin on functional brain architecture relative to placebo. Participants underwent imaging during dosing sessions with 15 mg psilocybin, 20mg 2C-B, or placebo (bittering agent). Acute rsfMRI was conducted approximately 110 min after each intervention administration. Doses of psilocybin and 2C-B were chosen based on psychotropic equivalence extrapolated from prior study data. This was originally corroborated by behavioural assessments of the sample (12), with additional descriptions of dose selection and drug purchase, and study provided therein.

### Altered state questionnaires

Following scanner exit, participants completed three retrospective self-report questionnaires. These included the Amsterdam Resting-State Questionnaire (ARSQ), administered approximately 130 minutes post-drug intake, which assessed ongoing cognition during rsfMRI (32). Additionally, participants completed the 5 Dimensions of Altered States Questionnaire (5D-ASC) and the Ego Dissolution Inventory (EDI), approximately 360 minutes post-drug intake, to evaluate subjective effects retrospectively (33, 34). Self-report visual analogue scales (VAS), measuring subjective high and effect intensity, were also completed immediately before and after the rsfMRI session (approximately 90 and 120 minutes post-drug intake). These assessments were used to reassess dose equivalency (12). For more details, see the Supplementary Materials.

### fMRI acquisition and preprocessing

Participants underwent one imaging session per acute dosing visit separated by a two-week washout, comprising structural imaging, magnetic resonance spectroscopy, task and rsfMRI. Across visits, imaging was performed within subjects at a fixed time of day to minimise diurnal effects (35). Images were acquired on a Siemens MAGNETOM 7T MRI scanner. Imaging parameters and procedures are detailed in the Supplementary Materials.

Neuroimaging data was preprocessed using fMRIPrep 1.1.147 (36). For details, see Supplementary Materials. Unsmoothed, preprocessed functional images were denoised using CONN 20.a(37). Following linear detrending, we implemented voxel-wise confound regression by regressing out (1) physiological noise from five aCompCor principal components (38), comprising white matter and cerebrospinal fluid signals (2) 12 motion parameters (3 translations, 3 rotations and first-order derivatives), (3) censoring regressors devised using the Artifact Detection Toolbox (ART), with a global signal threshold of z>3 and a framewise displacement (FD) threshold of >0.5 mm, (4) constant and (5) linear BOLD session signal trends to minimise the influence of slow trends and initial magnetisation non-steady-state transients. Time series were filtered using a 0.008–0.09 Hz band-pass filter. Our motion exclusion criterion comprised a mean FD greater than 0.5 mm and > 20% outlier scans in any scanning session. Quality control yielded 20 subjects, with no significant motion differences across conditions for either mean FD or censoring. To account for any possible residual motion artifacts, we replicated our findings using FD as a nuisance regressor. Secondary replication incorporating global signal regression (GSR) during denoising was also performed, given its influence on resting-state network composition (39). See Supplementary Materials for additional information.

Denoised BOLD timeseries were next parcellated into 232 regions, comprising 200 cortical regions of interest (ROIs) according to an augmented version of the functional parcellation of Schaefer et al. and 32 subcortical ROIs from the Tian atlas (40, 41). In addition, we repeated our primary analyses using a finer-grain 454 region parcellation (comprising Schaefer 400 and Tian 54, see Supplementary Materials).

### Functional organisation benchmarking

We derived spatiotemporal measures of resting-state network organization at both edge and regional levels to benchmark 2C-B’s effects. These measures have been extensively studied in the psychedelic literature (3, 4, 30). All connectomic measures were contextualized using the Yeo et al. 17 rsfMRI network classes (42).

Static functional connectivity (sFC) was computed by generating Pearson’s r between the timeseries of all pairwise combinations of 232 regions, with all r-values Fisher z-transformed. Global functional connectivity (gFC) serving as a proxy for node centrality, was calculated as the average Fisher z-transformed Pearson correlation from each region to all others.We focused on significant edges flagged by sFC statistics (43, 44).

To assess dynamic FC variability (dFC), we computed the variance of dynamic conditional correlations (DCC). DCC is a non-parametric model-based approach devised for assessing time-varying correlations (45). A framewise approach, it offers superior test-retest reliability compared to other dFC methods, which often face window artefacts and require arbitrary hyperparameters (46). Variance was calculated for all possible pairwise regional DCC combinations (47–49).

We selected and ranked spontaneous signal complexity measures according to previous replicability and association with psychedelic effects based on McCulloch et al.’s work, using the COPBET toolbox (31). Regional signal complexity was characterised using sample entropy (sampEn), defined as the negative logarithm of the conditional probability that two vectors of length m (set to 2) are dissimilar within a threshold distance r (set at 0.3 times the standard deviation of the regional timeseries) and that this dissimilarity persists when the vector length increases to m + 1 (50). From the same selection criteria, we also derived edgewise DCC Shannon entropy (dccEn), whole-brain Lempel-Ziv entropy (ZivEn) and path-length degree-distribution entropy (degreeEn). See Supplementary Materials for information on each measure.

### Pharmacological action mapping

Molecular densities were estimated from PET tracer studies for relevant serotonin receptors (5-HT_1A_, 5-HT_1B_, 5-HT_2A_) and monoamine transporters (serotonin reuptake [SERT], dopamine reuptake [DAT] and norepinephrine reuptake transporter [NET]). Targets were selected from literature demonstrating functional affinity of either 2C-B or psilocybin for these sites (14–16). Volumetric PET images were registered to standard space, averaged across participants within each study, and then parcellated into ROIs. Receptors having more than one mean image of the same tracer (5-HT_1B_) were combined using a weighted average. Regional values for cortical and subcortical areas were z-scored independently before their integration for analysis, considering established differences in radioligand uptake between these structures (51). For details on each PET dataset, see (52).

Dominance analysis was used to determine the relative importance of each PET density map in predicting regional contrast *t*-statistics for our primary imaging benchmarks (nodal sum sFC & dFC, sampEn). Dominance analysis aims to establish the relative significance (or “dominance”) of each independent variable in relation to the overall fit (adjusted R^2^) of the multiple linear regression. We also independently investigated the associative validity of the top topographical contributors for each metric and by performing spatial autocorrelation-preserving Pearson correlations. We generated spatially constrained nulls (10 000 iterations) using Moran Spectral Randomisation as implemented in the BrainSpace toolbox, a process first proposed in the ecology literature (53). This approach is best suited to assess the concordance between two spatial maps in which a geodesic projection cannot be ensured (e.g atlases comprising subcortical structures)(54). See Supplementary Materials for additional details.

### Experiential mapping

We assessed the behavioural relevance of “net” drug effects of both 2C-B and psilocybin on functional organisation by investigating the predictive value of regional multivariate coherence. Multivariate coherence, as presented herein, is defined as the Pearson correlation across measurements for a particular node. For each participant, primary nodal psychedelic effect measures (nodal sum sFC & dFC, sampEn) were *z*-scored across regions to control for inter-feature variability. Pearson correlations were then performed for each pair of regional feature vectors, forming an ROI x ROI matrix per participant, ultimately summed to generate a measure of nodal coherence strength per region. By performing a multivariate integration of measures of interest, this approach provides a simple regional summary of “net” drug effects while diminishing the number of potential multiple comparisons. This approach is comparable to generating annotation similarity networks (55).

We employed a multilevel partial-least-squares (PLS, accounting for inter-individual variability) analysis to assess the relationship between drug-induced effects on regional coherence and corresponding subjective effects (https://github.com/rmarkello/pyls). PLS is an unsupervised multivariate statistical approach that relates two feature matrices (X, Y) by estimating ther weighted linear combination to maximise their covariance. Here, X corresponds to multivariate coherence, with *n* being subjects and *r* being coherence change scores per condition (*X_nxr_*) and Y reflecting experiential measures, with *n* being subjects and *d* corresponding to change scores for each of our 6 retrospective dimensions (5D-ASC, EDI) per condition (*Y_nxd_*). This decomposition yields orthogonal latent variables (LVs) associated with a pattern of neural activity (i.e., multivariate coherence) and subjective effects (56). We assessed overall LV significance using permutation testing (10 000 permutations), and loading stability by generating bootstrap ratios (10 000 bootstraps). Overall model findings were also supported using 5-fold cross-validation, to derive out-of-sample correlation estimates. See Supplementary Materials.

### Statistics

For all analyses, the alpha criterion was set at *p <* l10.05 following appropriate multiple comparison correction.

A network-based statistic (NBS) approach (57) derived from the publicly available NBS toolbox was used to assess alterations to connectomic measures in a repeated measures design (see Supplementary Materials). Significant NBS (10 000 permutations) multiple-comparisons corrected edges were used as masks for post-hoc *t*-tests assessments. Changes to regional rsfMRI measures and behavioural outcomes were assessed by linear mixed effect models (LMEMs) with drug condition as a main fixed effect and participant as a random intercept. Follow-up assessments were performed with paired *t*-tests after Benjamini–Hochberg false discovery rate (FDR) correction across significant ROIs. For behavioural outcomes, Tukey’s method was applied.

## Results

We sought to define 2C-B’s effects on resting-state functional brain organisation and explore relative differences to psilocybin across a range of established psychedelic neuroimaging benchmarks.

### Administration of 2C-B and psilocybin yield comparable psychoactive effects

We first aimed to ensure psychotropic equivalence during resting-state fMRI acquisition (Figure 1, Table 1). Analyses revealed a significant drug effect on all VAS items immediately before (approximately 90 min post-administration) and after (approximatley 120 min post-administration) resting-state acquisition indicating comparable subjective effect intensity at the time of acquisition. The maximal drug effects (e_max_) for our real-time measures were equivalent for both 2C-B and psilocybin, consistent with previous findings (12). Retrospective dysphoric experiences related to changes in to selfhood (‘ego dissolution’), specifically anxious ego dissolution (AED), but not the Ego Dissolution Index (EDI), as well as the overall 5D-ASC (ASC) score, were significantly higher under psilocybin compared to both placebo and 2C-B.

**Figure 1.**
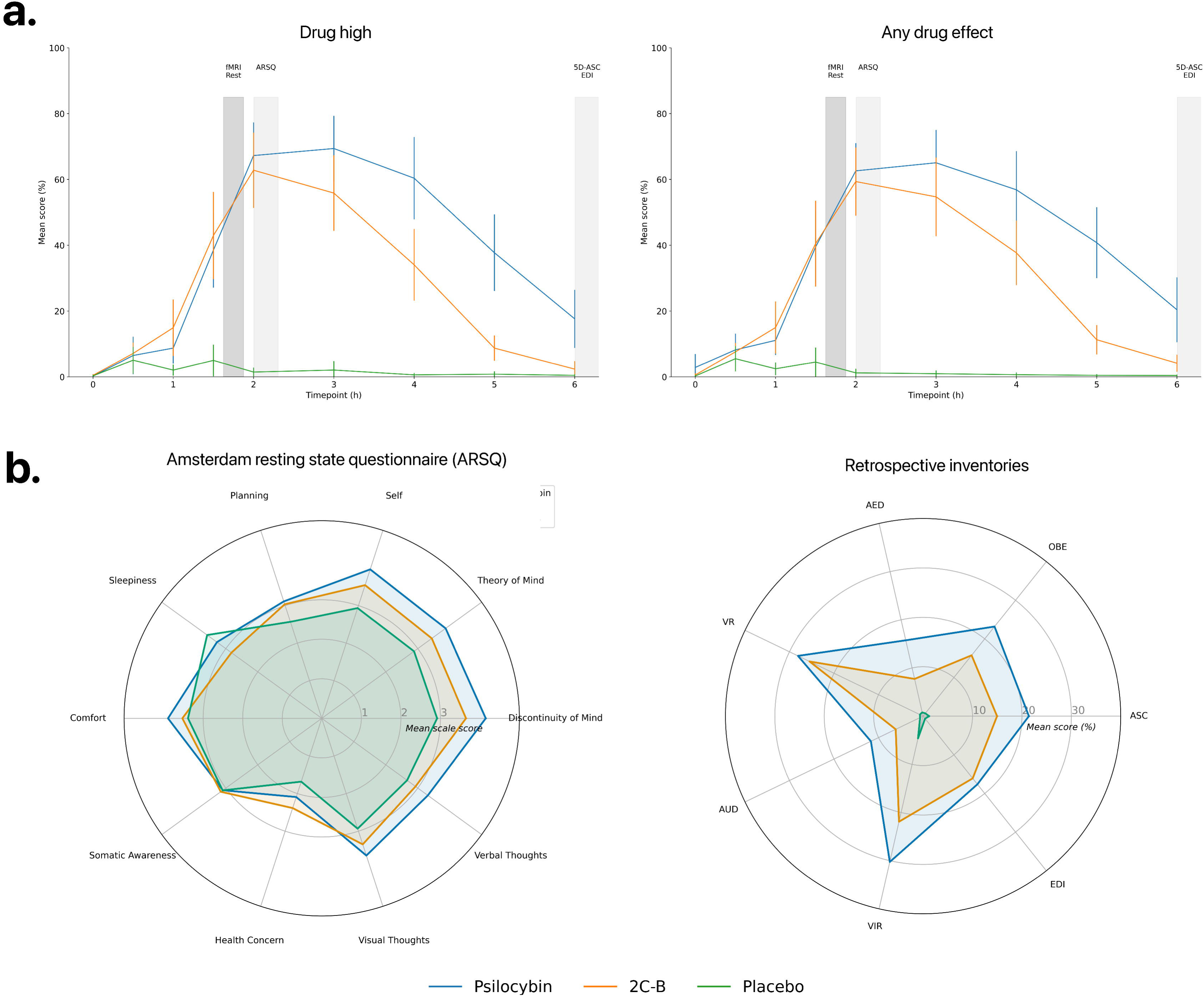
Self-reported measures of drug intensity and phenomenology. **A)** Time courses of each VAS measurement. Data are presented as meanlll±lllSEM (brackets) relative to scanning and phenomenological measures. **B)** Retrospective phenomenological measures (5D-ASC and EDI). Data are presented as means. Abbreviations: AED, anxious ego dissolution; OBE, oceanic boundlessness; ASC, altered state of consciousness; EDI, ego dissolution inventory; VIR, vigilance reduction; AUD, auditory alterations; VR, visual restructuralisation.

**Table 1.**
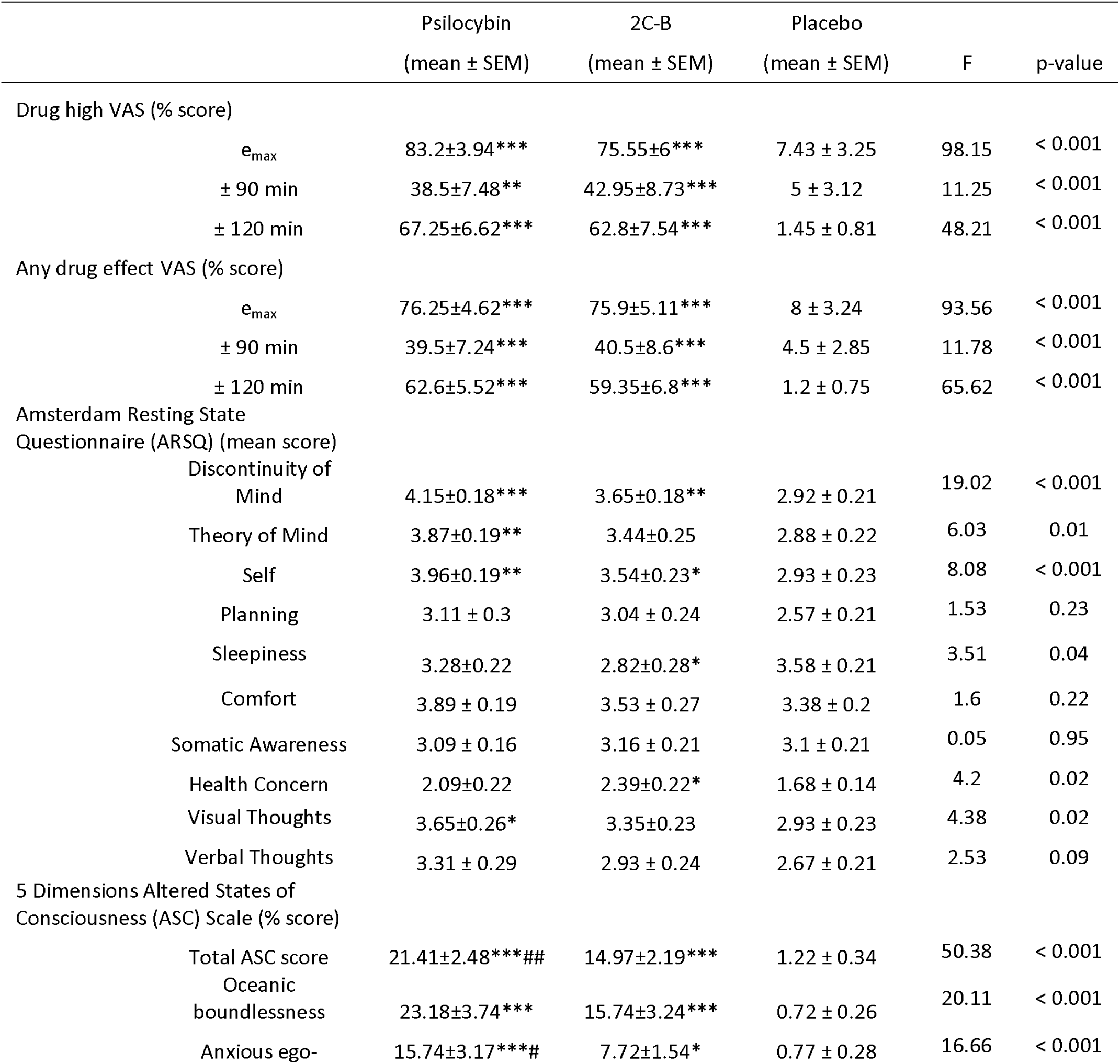
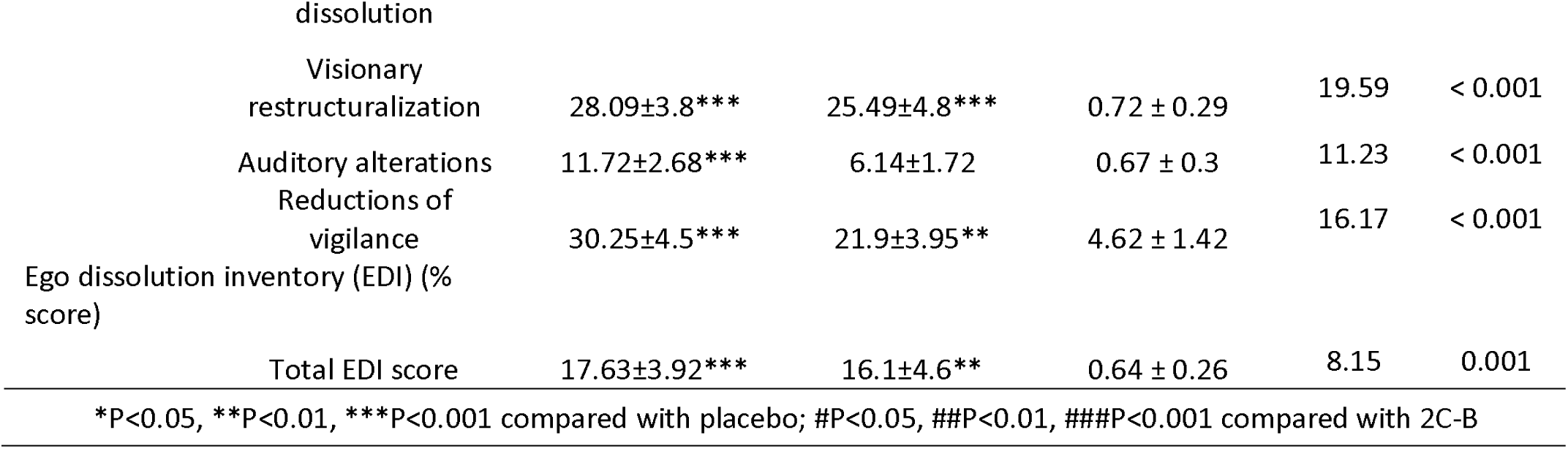
Comparison of subjective effects under 2C-B, psilocybin and placebo.

Following scanner exit, participants reported the phenomenological content of their resting-state mentation using the ARSQ. Analyses demonstrated a significant drug effect across 6/10 dimensions of resting-state cognition, with numerically large increases for both drugs relative to placebo in discontinuity of mind and self-referential thinking. However, neither compound exhibited significant differences in any ARSQ dimension during acquisition. Particularly, measures of in-scanner arousal (sleepiness) and comfort did not show compelling differences.

### Measures of brain organisation are differentially affected by 2C-B and psilocybin

Widespread changes in spatiotemporal brain organisation were observed under both drugs, consistent with prior reports of diminished unimodal-transmodal organization under psychedelics (see Figure 2). All outcomes were largely consistent after controlling for FD, across different parcellation schemes and global signal regression (see Supplementary).

**Figure 2.**
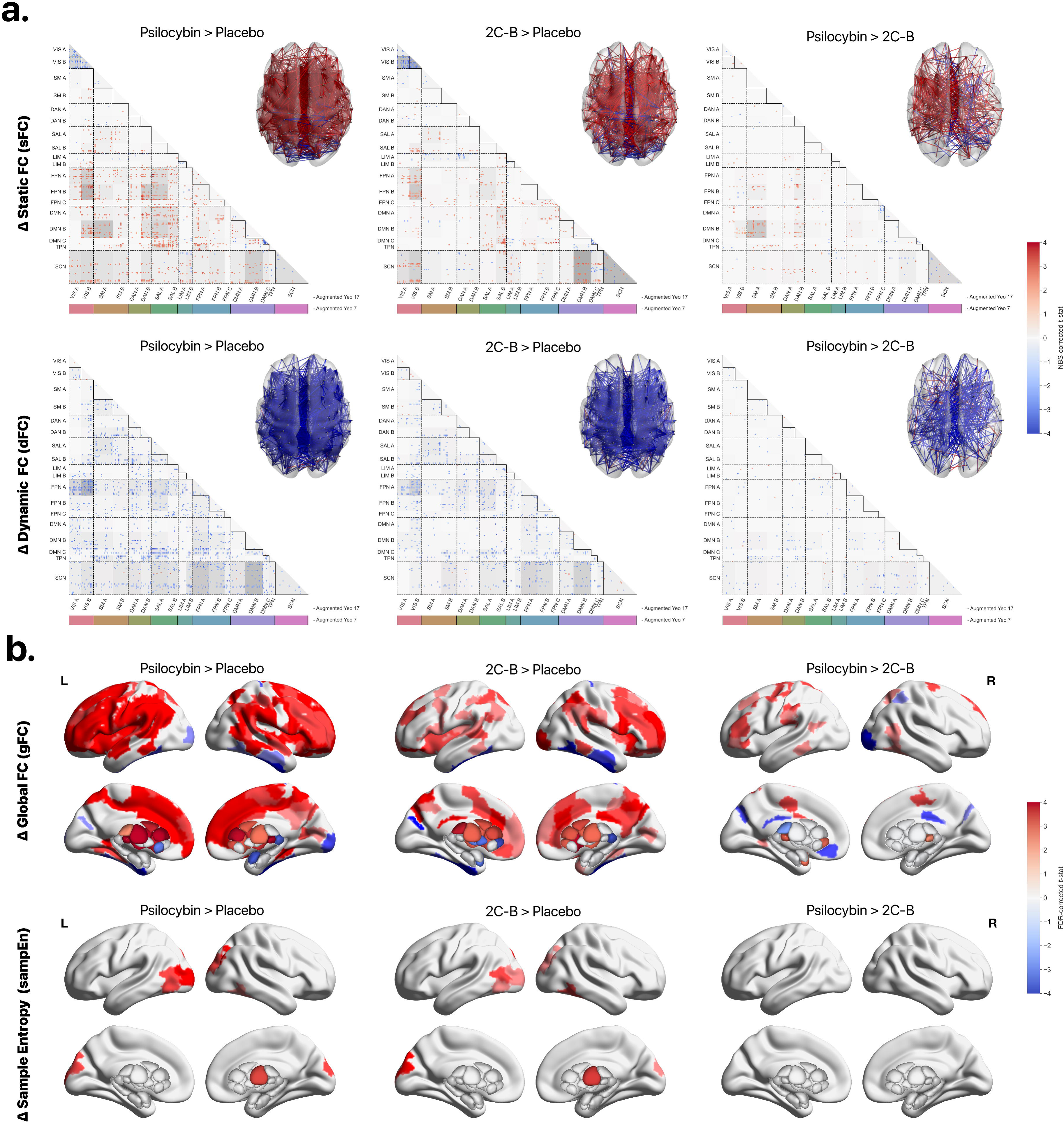
rsfMRI psychedelic effect benchmarks. **A)** Significant differences in drug-mediated static and dynamic FC across all contrasts. Cortical networks are based on the 17-network, 200-parcel Schaefer-Yeo parcellation (Schaefer et al. 2017) augmented with 32 subcortical regions from Tian et al. 2020. Red indicates drug-mediated increases in interregional FC, and blue indicates decreases. Grey hues represent the proportion of significant edges for a given Yeo 17 network pair relative to the total number of significant edges. Regions are ordered left and then right within each network. **B)** Significant differences in drug-mediated regional gFC (i.e., the average of a given seed region’s FC to the rest of the connectome) and sampEn (the complexity of a regional timeseries) across all contrasts. Abbreviations: VIS, visual; SMN, somatomotor network; DAN, dorsal attention network; SAL, salience network; LIM, limbic; FPN, frontoparietal network; DMN, default mode network; TPN (temporoparietal network); SCN (subcortical network).

For sFC, NBS identified widespread differences in sFC as indicated by a main effect of drug ( *p_NBS_* = 0.0421, 229 nodes, 1922 edges). Consistent with previous studies, psilocybin significantly reduced within-network connectivity for visual network A & B (VIS_A/B_) and the default mode network C (DMN_C_), compared to placebo. Network segregation, particularly across the frontoparietal network A & B (FPN_A/B_), salience network A & B (SAN_A/B_), and DMN_A/B/C_, was also significantly reduced. In addition, cortical-subcortical connectivity was broadly increased. 2C-B showed similar effects, with reductions in within-network sFC for VIS_B_ and DMN_C_, but showed converse increases in FPN_C_. Reductions in network segregation were focal, notably between FPN_A/B_ and VIS_A/B_, and SAL_B_ with transmodal networks. Subcortical-cortical sFC, especially involving DMN_B_ and VIS_B_, was significantly increased. Between-drug comparisons revealed that psilocybin had a generally greater reducing effect on between-network sFC relative to 2C-B, particularly for connectivity spanning the DMN_B_ and somatomotor network B (SMN_B_), whereas 2C-B showed greater increases in DMN_A_-FPN_C_ connectivity. These results align with a predominantly cortical effect, highlighting the significance of transmodal cortices.

For dFC, NBS analyses also revealed a widespread effect of drug (*p_NBS_*= 0.0135, 231 nodes, 1755 edges). Both psilocybin and 2C-B significantly induced strong, nonspecific whole-brain reductions in between-network dFC variance compared to placebo, with the greatest decreases observed in subcortical-cortical (DMN_B_, FPN_A/B_) and VIS_A_-FPN_A_ connectivity variance. No clear within-network effects were visible under either compound. Post-hoc comparisons across compounds revealed overall spatially distributed greater reductions in dFC under psilocybin relative to 2C-B for between-network dFC. Edgewise complexity (dccEn) did not exhibit significant changes (*p_NBS_* > 0.1).

In parallel, we calculated regional measures of gFC and several complexity measures (Figure 2B, Supplementary). gFC analyses showed significant increases in mean FC across most regions for both substances compared to placebo. Under psilocybin, notable increases were seen in the left dorsolateral and bilateral ventral prefrontal cortex, inferior frontal gyrus, and left ventral posterior thalamus, with decreases in the right temporal pole, medial amygdala, dorsal posterior thalamus, and occipital lobe. For 2C-B, the strongest gFC increases occurred in the right precuneus, lateral prefrontal cortex, left posterior cingulate cortex, and right insula, with reductions in the bilateral temporal poles, right dorsal posterior thalamus, and left nucleus accumbens. Between drugs, psilocybin generally led to greater gFC increases compared to 2C-B, particularly in the left postcentral gyrus, medial cingulate cortex, ventral posterior thalamus, and frontal eye fields. In contrast, 2C-B showed greater increases in the right extrastriate visual cortex, bilateral posterior cingulate, medial prefrontal cortex, and left dorsal anterior thalamus. Controlling for motion, GSR exhibited drug-dependent effects (see Supplementary Materials).

Analyses of regional complexity using sampEn revealed comparable effects for 2C-B and psilocybin, indicating an increase in time series complexity. Specifically, both compounds significantly increased sampEn in the bilateral extrastriate/primary visual cortex and the anterior ventral thalamus, with no significant differences observed between the compounds. Whole-brain signal complexity (zivEn but not degreeEn) was also significantly increased relative to placebo, with no significant differences between compounds (see Supplementary Materials).

### Changes to functional brain organisation follow pharmacology

Through dominance analysis, we compared the strength of primary regional metrics’ associations with pharmacological receptor and transporter maps (see Figure 3A). Separate models were generated for each primary outcome measure (sFC, dFC and sampEn nodal sum *t*-stats) per contrast and were supported by independent correlations (Figure 3B). Across 54 possible models, we found that for both 2C-B and psilocybin, dFC changes from placebo were best explained (Psilocybin: 56.89%, 2C-B: 64.38 %) by an inverse relationship with 5-HT_2A_ receptor density indicating that dFC variance decreases relative to placebo, 5-HT_2A_ spatial density increases. Similarly, sampEn was best explained for each drug (Psilocybin: 48.26%, 2C-B: 45.40 %) by an inverse relationship with 5-HT_1A_ spatial density, suggesting increased signal complexity relative to placebo correlates with lower 5-HT_1A_ levels. For 5-HT_1B_, there was a significant positive relationship with sFC under psilocybin (41.89%), suggesting greater increases in sFC for regions of high 5-HT_1B_ density.

**Figure 3.**
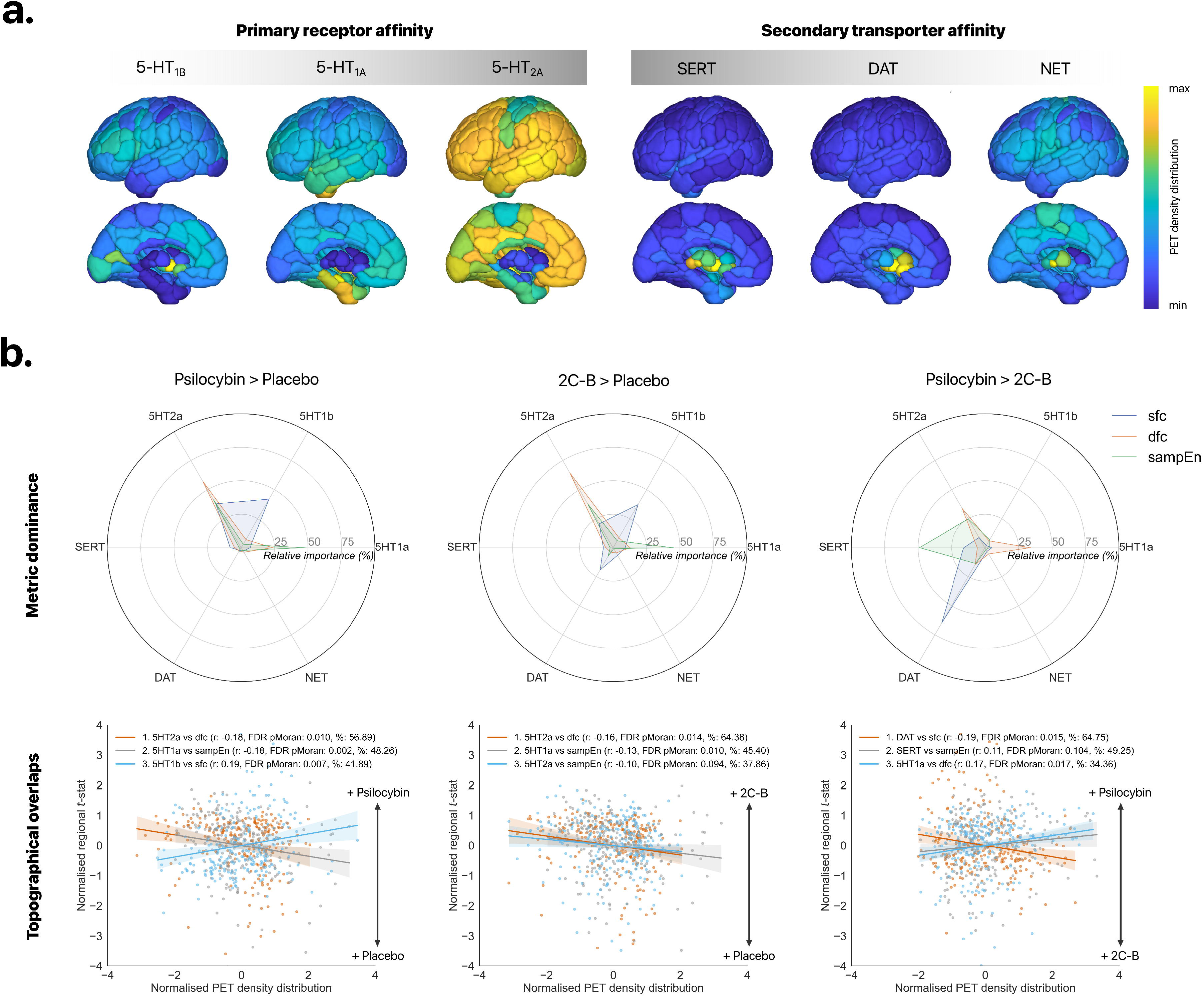
Pharmacology-guided mapping of rsfMRI outcomes. **A)** PET molecular density distributions of primary and secondary drug affinities are displayed. Values are expressed as raw expression values prior to z-scoring. **B)** Top: Dominance analysis for each contrast and measure of interest. Dominance analysis distributes the fit of the model across input receptor maps such that the contribution of each receptor can be assessed and compared to other input maps. The percent contribution of each input map is defined as the map’s dominance normalized by the total fit (adj2) of the model. Bottom: ranked scatter plots of top predictive receptor-outcome measure pairs. Contrast *t*-statistics are plotted relative to z-scored receptor distributions.

Importantly, when assessing relative differences across drugs, DAT density was a strong determinant of topological differences in sFC between 2C-B and psilocybin (65.75%). Specifically, areas with greater sFC under 2C-B relative to psilocybin correlated with higher DAT density and vice versa. Whereas SERT was found to be a strong contributor to sampEn between-drug differences (49.25%) independent correlations revealed a non-significant, positive relationship. Lastly, 5-HT_1A_ was found to also significantly contribute to between-drug differences (34.36%): in regions with lower 5-HT_1A_ density, psilocybin tended to exert a stronger effect on reducing dFC variance compared to 2C-B ( and vice versa).]

### Multivariate neural effects show distinct associations with subjective experience

We lastly sought to understand how net effects on functional brain organisation may relate to a participant’s subjective experience. To do so, we generated multivariate coherence change scores per region and drug, summarising the nonlinear effects of each compound across largely uncorrelated measures (see Supplementary Materials). As depicted in Figure 4A, multivariate coherence followed a unimodal-transmodal topographical organisation. Primary networks (e.g., VIS, SM) closely coupled to sensory input, showed high multivariate coherence and responded similarly across measures, while more flexible heteromodal networks exhibited lower coherence, responding asynchronously. Levene’s testing indicated the variance of regional multivariate coherence scores differed across conditions (W = 22 p < 0.0001).

**Figure 4.**
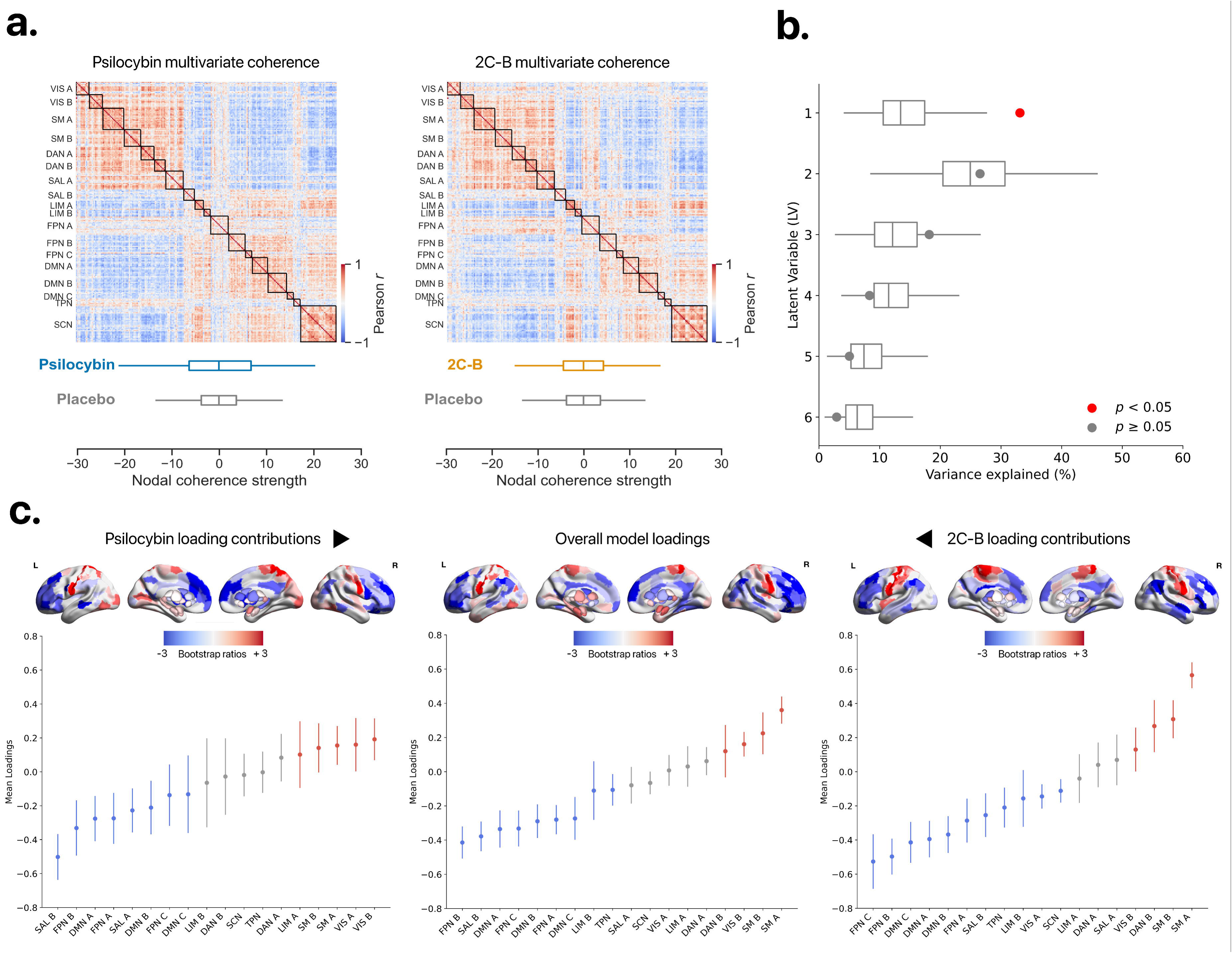
Multivariate contributions to behavioural prediction. **A)** Top: multivariate coherence matrices per drug. For a given pair of brain regions, metrics are correlated (Pearson *r*) to yield a measure of net drug effects per corresponding edge. Bottom: distribution of regional multivariate coherence scores (nodal sum of each matrix) per condition. **B)** The first LV (PLS1) was statistically significant, concerning null model permutation testing in grey (10,000 permutations, depicted as grey boxplots). The absolute model explained variances are represented as dots. **C)** Regional model loading contributions for PLS 1. Warm values indicate a positive contribution to the LV whereas cold values represent a negative contribution. Top: Brain renders represent derived loading bootstrap ratios (calculated across 10,000 iterations). Bottom: corresponding raw model loadings are aggregated below into their respective Yeo-17 networks, ranked according to their means and plotted with their standard error of means (1.5). Network loading sign directions follow bootstrap ratio scores. Abbreviations: VIS, visual; SMN, somatomotor network; DAN, dorsal attention network; SAL, salience network; LIM, limbic; FPN, frontoparietal network; DMN, default mode network; TPN (temporoparietal network); SCN (subcortical network).

We next applied a multilevel PLS analysis to identify a multivariate mapping of maximum covariance between drug-induced effects on brain organisation and subjective effects. Permutation testing revealed a statistically significant latent variable (PLS1), explaining 33.13 % of total cross-drug experiential scores (*p_perm_* = 0.0291). Across contrasts, regional coherence PLS1 scores were positively associated with experiential PLS1 scores (overall: *r*_in-sample_ = 0.40, *r*_out--sample_ = 0.32, *p_FDR_* = 0.0151; Psilocybin: *r*_in-sample_ = 0.79, *p_FDR_* = 0.0001, 2C-B: *r*_in-sample_ = 0.53, *p_FDR_* = 0.0151), suggesting PLS1 captured a pattern in multivariate coherence that is positively associated to subjective effects.

To evaluate spatial patterns of covariance relevant to each drug’s effects, we extracted PLS1 regional loadings per condition after bootstrapping (Figure. 4C). Regions with positive loadings in PLS1 indicate that greater coherence following drug intake is linked to stronger subjective effects, while regions with negative loadings suggest that greater coherence in those areas is linked to weaker subjective effects and vice versa. Under psilocybin, the strongest negative loadings were in the prefrontal cortex (PFC), including SAL_B_, DMN_A_, and FPN_B_ (e.g., bilateral ventral, dorsal, and lateral PFC), while positive loadings appeared in primary sensory regions (e.g., striate/extrastriate cortex, precentral gyrus). 2C-B showed a similar spatial pattern, with subtle regional differences. Negative loadings were strongest in regions associated with FPN_B/C_ and DMN_C_ (e.g., bilateral inferior parietal lobe, dorsal and lateral PFC), and positive loadings appeared in areas linked to SM_A/B_ and DAN_B_ (e.g., primary/secondary motor cortex, postcentral gyrus).

Overall, PLS1 suggests that a transmodal cortical gradient best characterises psychedelic effects. Cortical PLS1 bootstrap ratios correlated with the primary gradient of cortical FC (58) and the sensorimotor-associative (S-A) axis of brain organisation (59) following spin permutation tests (FC gradient: *r* = 0.50, *p*_FDRspin_ < 0.0001; S-A axis: *r* = −0.51, *p*_FDRspin_ < 0.0001, see Supplementary). Across psychedelics, regional neurobehavioral markers are thus marked by reduced multivariate coherence in transmodal networks (including regions such as the PFC and temporoparietal junction) and increased coherence in sensorimotor cortices.

## Discussion

This study aimed to characterise the neural correlates of 2C-B using an integrative approach that combines spatiotemporal measures of functional organisation, molecular pharmacology, and subjective experience. Consistent with our hypothesis, we found that 2C-B exhibits empirical properties similar to classical psychedelics, including network desegregation and increased spontaneous BOLD complexity, irrespective of variations in rsfMRI thought content or effect intensity. By examining functional (sub)networks, we identified coupling changes to subcortical and prefrontal regions as well as transmodal network desegregation across compounds. Furthermore, we confirm that regional differences between 2C-B and psilocybin effects are associated with reported pharmacodynamic differences in 5-HT_1A_ and DAT affinity, with serotonergic receptors sharing a common role in defining functional dynamics. Characterising nonlinear psychedelic effects, we isolated neurobehavioral markers for each compound, contributing to a net destabilisation of prefrontal cortices. Our findings suggest that 2C-B is a valuable tool for psychedelic neuroscience, while enhancing our understanding of the differing neurobiological mechanisms of psychedelics.

One of the strongest findings in psychedelic research is the reduction of intranetwork sFC for the VIS and DMN, often termed network ‘disintegration’. This effect is observed across various psychedelics and entactogens, such as lysergic acid diethylamide (LSD), ayahuasca, dimethyltryptamine (DMT), psilocybin, and MDMA (43, 60–63). Given 2C-B’s compatible phenomenology, its similarity here is expected. Previous studies (30) have often used broad network definitions (e.g. Yeo 7) or methods like independent components analysis (ICA), which may overestimate intranetwork effects by examining coherent neural activity across extended parts of the brain. Here for example, while within-network sFC was broadly reduced for the VIS under 2C-B and psilocybin, only the DMN_C_ (comprising the retrosplenial cortex, parahippocampus, and anterior hippocampus) showed reduced sFC. This supports the need for finer (sub)network definition, as highlighted by a recent precision hypersampling approach demonstrating that psilocybin’s enduring effects are localised to altered hippocampal-cortical connectivity (64).

It is theorised that the degree of whole-brain functional integration underlies the depth of the psychedelic experience (65). Psychedelics with rich phenomenological profiles, such as DMT show a substantial compression of functional cortical organisation (43), whereas minimally “disruptive” compounds such as MDMA or dexamphetamine, result in less extensive changes (44, 66). In our study, compared to psilocybin, 2C-B demonstrated less extensive unimodal-transmodal internetwork sFC (e.g., DMN_B_, SM_A_) and subcortical-cortical connectivity, but exhibited enhanced transmodal connectivity (e.g. DMN_C_-FPN_A_). The use of gFC revealed marked increases in connectivity within thalamic nuclei involved in sensorimotor gating (67) and flexible network reconfiguration (68) and substantial portions of the neocortex for both compounds. Interestingly, 2C-B was associated with marked increases in gFC and sFC compared to psilocybin for areas such as the mPFC (subdivisions of corresponding to the FPN and DMN), which impacted a broader extent of the neocortex. Sparse increases in FPN sFC after MDMA (63) and reductions in centrality after SSRIs have been observed (69). Whereas our findings follow those of groups showing increased transmodal and reduced unimodal gFC (43, 61, 70) diverging findings have been identified (71, 72). Further work is needed to reconcile these differences given that incorporating GSR seemingly taps into latent pharmacological properties (see Supplementary).

Temporal assessments of brain organisation across substances revealed significant reductions in dFC variance across the brain and increased spontaneous signal complexity. Notably, 2C-B yielded less widespread reductions in dFC variance compared to psilocybin while increases in regional entropy were consistent between the two compounds, indicating that the between-drug differences in the minimisation of drastic connectivity changes are independent of an increased prevalence of unique signalling motifs. This interpretation may be mutually equivalent to results from cluster-based dynamical analyses, showing greater occurrence rates of hyperconnected FC states under psychedelics (73, 74). Acute reductions to dFC variance parallel to numerically increased entropy, have also been observed after inhalation of the dissociative hallucinogen Salvia divinorum (48). Doss and colleagues however report subacute increases in dFC variance following psilocybin-assisted therapy for depression, which they refer to as a marker of increased ‘neural flexibility’ following treatment - otherwise absent in healthy volunteers (47, 49). Further replication is needed to validate the concept of neural flexibility in the acute psychedelic state and its relationship to longer-term effects.

Whole-brain complexity, which refers to the irreducibility of brain activity from functional integration and differentiation, is considered a hallmark of the phenomenological richness of psychedelics (75). Eliciting similar behavioural effects, 2C-B also produced comparable increases in complexity to psilocybin. It remains to be studied however whether this is a feature inherent to 5-HT_2A_ agonism, since MDMA or amphetamines have yet to be assessed. We particularly observed increased complexity in thalamic nuclei under psychedelics, aligning with the ‘thalamic gating’ model of psychedelic effects that suggest psychedelics disinhibit thalamic connections, allowing more sensory information to reach the neocortex (76). It should be noted that sparse regional elevations in entropy were identified, with no significant changes in edge-based complexity. This may be reflective of our choice of eyes-open rsfMRI to minimise arousal differences as it provides a more ‘grounded’ psychological state, akin to task-based approaches where attention is directed towards external stimuli rather than internal psychological processes. Studies indicate that complexity scales with cognitive load (77) and external stimulation under psilocybin (64), with no significant changes observed during eyes-open resting EEG (78). A more definitive consensus on the applicability of different measures of complexity could thus be gleamed from block designs increasing in task demand (79).

For the development of novel compounds with refined affinities in psychedelic-based pharmacotherapy, it is crucial to understand how molecular pharmacodynamics translate into cortical markers of therapeutic benefit. Extending a body of work relating 5-HT_2A_ agonism to altered regional dynamics (24–26) our ranked modelling approach identifies regional 5-HT_2A_ expression to most strongly align with greater reductions in dFC across psychedelics relative to placebo. Concurrently, regional 5-HT_1A_ density was found to correlate with changes in sampEn relative to placebo and dFC differences under psilocybin relative to 2C-B. Coexpressed, 5-HT_1A_ autoreceptors act in opponency to 5-HT_2A_ receptors to dampen the spiking output of cortical pyramidal neurons, attenuating changes to cortical excitability (80) and perhaps by extension, signal complexity. Recent theoretical modelling of cortical dynamics has proposed that between-compound differences in 5-HT_1A_ agonism may mediate psychological tolerability (81). Considered a target for antidepressant treatment (82), 5HT_1B_ emerged as a novel marker for psilocybin’s effects on sFC. 5-HT_1B_ autoreceptors are expressed on cortical serotonergic neurons as well as the raphe and basal ganglia (83, 84), reducing synaptic 5-HT release via negative feedback mechanisms (85). Furthermore, 5-HT_1B_ receptors act as heteroreceptors regulating the release of other neurotransmitters, including GABA and dopamine (82). Across compounds, we also found that higher DAT density was associated with increased sFC under 2C-B relative to psilocybin. This aligns with 2C-B’s secondary monoaminergic binding affinity and preclinical work showing increased striatal dopamine and reduced EEG coherence post-administration (86). With 2C-B reported to yield less discontinuity of mind compared to other psychedelics (12, 87), it is worth speculating whether its dopaminergic effects may contribute to this observation. The degree of cognitive lability under psychedelics has been described to be reliant on cognitive control mechanisms which mediate the balance between “stable” and “divergent” thinking via the optimisation of prefrontal-striatal dopaminergic transmission (88). Exploring ways to operationalise dopaminergic tone to either enhance cognitive flexibility or reduce impairment may provide new options for clinical populations characterised by cognitive rigidity.

Our analyses also sought to distinguish behavioural correlates of “net” drug effects on brain function. Here, we found that asynchronicity in transmodal cortices (i.e. reduced coherence), which are involved in high-level cognitive processing (58, 59, 89) serves as a useful neurobehavioral marker of psychedelic effects. Additionally, our results indicate that the behavioural impacts of each drug are particularly contingent on the destabilisation of specific subnetworks within the DMN, FPN and SAL – together forming the triple network of psychopathology posited to underlie the symptomatology of mood disorders (90). Given that interindividual variability in complex cognition under both normative (91) and psychedelic states (92, 93) lies in these evolutionarily recent brain regions, clarifying compound-specific regional markers may provide future avenues for personalising psychedelic-assisted therapy. Theoretically, compound selection based on their ability to maximise neural therapeutic effects could be further steered by intervention-based modelling approaches to identify cortical perturbation points (94, 95). It should be noted that psychedelics impair some aspects of memory recollection in humans (96) which may impact the fidelity of post-hoc subjective ratings and how they correlate with acute, objective brain measures. Analysing task-based fMRI, we will next seek to characterise how nonlinear dynamics may map onto relevant real-time cognitive processes such as emotional decision-making.

The present findings are not without limitations. Firstly, a larger sample size would have enabled a more robust detection of between-drug effects and behavioural correlates. Efforts are ongoing to integrate our results into a multi-group consortium characterising psychedelics. Secondly, we acknowledge there is a broader range molecular targets, as well as relevant complexity measures beyond those validated by McCullough et al. This is pertinent given the current absence of *in vivo* 2C-B receptor occupancy data and the nonspecificity of 5-HT_2A_ PET tracers (CIMBI36) for 5-HT_2A/2C_ receptors due to high sequence homology (7, 97). Thirdly, the timing of our subjective intensity queries makes it difficult to ascertain whether the resting-state fMRI acquisition aligned with the late onset phase or peak psychoactive effects. The magnitude of observed effects may thus differ according to the asssessed temporal phase (71). Finally, an empirical question persists: how does one best compare ‘apples and oranges’ in psychedelic neuroscience? The subjective nature and variable duration of these experiences, coupled with a complex underlying polypharmacology beyond 5-HT_2A_ agonism, raises significant challenges for defining *in vivo* equivalences. As demonstrated here and by others (44), a perhaps ecologically sound approach involves matching single doses across relevant behavioral or cardiovascular outcomes. Dose-ascending pharmaco-imaging studies with numerous active controls are likely to be costly and burdensome for participants while further accentuating the risk of order effects due to the persisting effects of psychedelics (98). Moving forward, intelligent comparative study design will be key to evaluating the relative benefits and drawbacks of different psychedelics for therapy.

In conclusion, this 7T rsfMRI study demonstrates that 2C-B induces significant multivariate disruptions in the brain’s spatiotemporal organisation, expanding our understanding of classic psychedelics. A direct within-subject fMRI comparison of two 5-HT_2A_ agonists, our findings show secondary pharmacodynamics may actively shape psychedelic effects and highlight new ventures for psychedelic drug development.

## Data availability

All code used for analysis and derivatives are to be made available at the following link: (https://github.com/PabloMallaroni/project_cb_rsfmri). Raw data and accompanying covariates can be made available to qualified research institutions upon request to J.G.R and a data use agreement executed with Maastricht University.

## Author contributions

Conceptualisation - P.M; Investigation - P.M, N.L.M, J.G.R; Project Administration - P.M, N.L.M, J.G.R; Formal analysis - P.M; Methodology - P.M, P.S; Software - P.M, P.S; Validation - P.S; Writing – Original Draft, P.M, Writing – Review & Editing; P.M, P.S, N.L.M, T.S, J.G.R.; Visualization - P.M; Funding Acquisition - J.G.R; Supervision - J.G.R, N.L.M.

## Acknowledgements

We thank Manoj K. Doss for discussions pertaining to DCC interpretation, Drummond E. McCulloch for the implementation, maintenance of and feedback on the COPBET toolbox, Emma de Brabander for assistance in data collection, Enrico Amico for advice on statistical approaches. We thank our volunteers for participating in our study.

## Conflict of interest

Johannes G. Ramaekers is a scientific consultant to GH Research who have no involvement in the preparation or conception of this manuscript or related data. All other authors have no relevant conflicts of interest to declare.

## Funding

Johannes G. Ramaekers acknowledges financial support from the Dutch Research Council (NWO, grant number 406.18. GO.019).

## References

1. Daws RE, Timmermann C, Giribaldi B, Sexton JD, Wall MB, Erritzoe D, et al. Increased global integration in the brain after psilocybin therapy for depression. Nature Medicine. 2022;28(4):844–51.

2. Deco G, Sanz Perl Y, Johnson S, Bourke N, Carhart-Harris RL, Kringelbach ML. Different hierarchical reconfigurations in the brain by psilocybin and escitalopram for depression. Nature Mental Health. 2024;2(9):1096–110.

3. Linguiti S, Vogel JW, Sydnor VJ, Pines A, Wellman N, Basbaum A, et al. Functional imaging studies of acute administration of classic psychedelics, ketamine, and MDMA: Methodological limitations and convergent results. Neuroscience & Biobehavioral Reviews. 2023;154:105421.

4. Girn M, Rosas FE, Daws RE, Gallen CL, Gazzaley A, Carhart-Harris RL. A complex systems perspective on psychedelic brain action. Trends in Cognitive Sciences. 2023;27(5):433–45.

5. Kwan AC, Olson DE, Preller KH, Roth BL. The neural basis of psychedelic action. Nature Neuroscience. 2022;25(11):1407–19.

6. Wall MB, Harding R, Zafar R, Rabiner EA, Nutt DJ, Erritzoe D. Neuroimaging in psychedelic drug development: past, present, and future. Mol Psychiatry. 2023;28(9):3573–80.

7. McClure-Begley TD, Roth BL. The promises and perils of psychedelic pharmacology for psychiatry. Nature Reviews Drug Discovery. 2022;21(6):463–73.

8. E-Wen McCulloch D, Liechti ME, Kuypers KPC, Nutt D, Lundberg J, Stenbæk DS, et al. Knowledge gaps in psychedelic medicalisation: Clinical studies and regulatory aspects. Neuroscience Applied. 2024;3:103938.

9. Mallaroni P, Mason NL, Vinckenbosch FRJ, Ramaekers JG. The use patterns of novel psychedelics: experiential fingerprints of substituted phenethylamines, tryptamines and lysergamides. Psychopharmacology. 2022;239(6):1783–96.

10. Sessa B, Fischer FM. Underground MDMA-, LSD- and 2-CB-assisted individual and group psychotherapy in Zurich: Outcomes, implications and commentary. Drug Science, Policy and Law. 2015;2:2050324515578080.

11. Doss MK, Mallaroni P, Mason NL, Ramaekers JG. Psilocybin and 2C-B at Encoding Distort Episodic Familiarity. Biol Psychiatry Cogn Neurosci Neuroimaging. 2024.

12. Mallaroni P, Mason NL, Reckweg JT, Paci R, Ritscher S, Toennes SW, et al. Assessment of the Acute Effects of 2C-B vs. Psilocybin on Subjective Experience, Mood, and Cognition. Clinical Pharmacology & Therapeutics. 2023;114(2):423–33.

13. Marcher-Rørsted E, Halberstadt AL, Klein AK, Chatha M, Jademyr S, Jensen AA, Kristensen JL. Investigation of the 2, 5-dimethoxy motif in phenethylamine serotonin 2A receptor agonists. ACS Chemical Neuroscience. 2020;11(9):1238–44.

14. Rickli A, Moning OD, Hoener MC, Liechti ME. Receptor interaction profiles of novel psychoactive tryptamines compared with classic hallucinogens. Eur Neuropsychopharmacol. 2016;26(8):1327–37.

15. Rickli A, Luethi D, Reinisch J, Buchy D, Hoener MC, Liechti ME. Receptor interaction profiles of novel N-2-methoxybenzyl (NBOMe) derivatives of 2,5-dimethoxy-substituted phenethylamines (2C drugs). Neuropharmacology. 2015;99:546–53.

16. Ray TS. Psychedelics and the human receptorome. PLoS One. 2010;5(2):e9019.

17. Zwartsen A, Verboven AHA, van Kleef RGDM, Wijnolts FMJ, Westerink RHS, Hondebrink L. Measuring inhibition of monoamine reuptake transporters by new psychoactive substances (NPS) in real-time using a high-throughput, fluorescence-based assay. Toxicology in Vitro. 2017;45:60–71.

18. Liechti ME, Baumann C, Gamma A, Vollenweider FX. Acute Psychological Effects of 3,4-Methylenedioxymethamphetamine (MDMA, “Ecstasy”) are Attenuated by the Serotonin Uptake Inhibitor Citalopram. Neuropsychopharmacology. 2000;22(5):513–21.

19. Becker AM, Holze F, Grandinetti T, Klaiber A, Toedtli VE, Kolaczynska KE, et al. Acute Effects of Psilocybin After Escitalopram or Placebo Pretreatment in a Randomized, Double-Blind, Placebo-Controlled, Crossover Study in Healthy Subjects. Clin Pharmacol Ther. 2022;111(4):886–95.

20. Robbins TW, Arnsten A. The neuropsychopharmacology of fronto-executive function: monoaminergic modulation. Annual review of neuroscience. 2009;32(1):267–87.

21. Avery MC, Krichmar JL. Neuromodulatory systems and their interactions: a review of models, theories, and experiments. Frontiers in neural circuits. 2017;11:108.

22. Shine JM, Aburn MJ, Breakspear M, Poldrack RA. The modulation of neural gain facilitates a transition between functional segregation and integration in the brain. Elife. 2018;7:e31130.

23. Shine JM, Breakspear M, Bell PT, Ehgoetz Martens KA, Shine R, Koyejo O, et al. Human cognition involves the dynamic integration of neural activity and neuromodulatory systems. Nature Neuroscience. 2019;22(2):289–96.

24. Deco G, Cruzat J, Cabral J, Knudsen GM, Carhart-Harris RL, Whybrow PC, et al. Whole-Brain Multimodal Neuroimaging Model Using Serotonin Receptor Maps Explains Non-linear Functional Effects of LSD. Curr Biol. 2018;28(19):3065–74.e6.

25. Singleton SP, Luppi AI, Carhart-Harris RL, Cruzat J, Roseman L, Nutt DJ, et al. Receptor-informed network control theory links LSD and psilocybin to a flattening of the brain’s control energy landscape. Nature Communications. 2022;13(1):5812.

26. Singleton SP, Timmermann C, Luppi AI, Eckernäs E, Roseman L, Carhart-Harris RL, Kuceyeski A. Time-resolved network control analysis links reduced control energy under DMT with the serotonin 2a receptor, signal diversity, and subjective experience. bioRxiv. 2023.

27. Pokorny T, Preller KH, Kraehenmann R, Vollenweider FX. Modulatory effect of the 5-HT1A agonist buspirone and the mixed non-hallucinogenic 5-HT1A/2A agonist ergotamine on psilocybin-induced psychedelic experience. European Neuropsychopharmacology. 2016;26(4):756–66.

28. Ballentine G, Friedman SF, Bzdok D. Trips and neurotransmitters: Discovering principled patterns across 6850 hallucinogenic experiences. Science Advances. 2022;8(11):eabl6989.

29. Luppi AI, Hansen JY, Adapa R, Carhart-Harris RL, Roseman L, Timmermann C, et al. In vivo mapping of pharmacologically induced functional reorganization onto the human brain’s neurotransmitter landscape. Science Advances. 2023;9(24):eadf8332.

30. McCulloch DEW, Knudsen GM, Barrett FS, Doss MK, Carhart-Harris RL, Rosas FE, et al. Psychedelic resting-state neuroimaging: A review and perspective on balancing replication and novel analyses. Neuroscience & Biobehavioral Reviews. 2022;138:104689.

31. McCulloch DE-W, Olsen AS, Ozenne B, Stenbæk DS, Armand S, Madsen MK, et al. Navigating the chaos of psychedelic neuroimaging: A multi-metric evaluation of acute psilocybin effects on brain entropy. medRxiv. 2023:2023.07.03.23292164.

32. Diaz BA, Van Der Sluis S, Moens S, Benjamins JS, Migliorati F, Stoffers D, et al. The Amsterdam Resting-State Questionnaire reveals multiple phenotypes of resting-state cognition. Front Hum Neurosci. 2013;7:446.

33. Studerus E, Gamma A, Vollenweider FX. Psychometric evaluation of the altered states of consciousness rating scale (OAV). PLoS One. 2010;5(8):e12412.

34. Nour MM, Evans L, Nutt D, Carhart-Harris RL. Ego-dissolution and psychedelics: validation of the ego-dissolution inventory (EDI). Frontiers in human neuroscience. 2016:269.

35. Shannon BJ, Dosenbach RA, Su Y, Vlassenko AG, Larson-Prior LJ, Nolan TS, et al. Morning-evening variation in human brain metabolism and memory circuits. Journal of Neurophysiology. 2013;109(5):1444–56.

36. Esteban O, Markiewicz CJ, Blair RW, Moodie CA, Isik AI, Erramuzpe A, et al. fMRIPrep: a robust preprocessing pipeline for functional MRI. Nature Methods. 2019;16(1):111–6.

37. Whitfield-Gabrieli S, Nieto-Castanon A. Conn: a functional connectivity toolbox for correlated and anticorrelated brain networks. Brain connectivity. 2012;2(3):125–41.

38. Behzadi Y, Restom K, Liau J, Liu TT. A component based noise correction method (CompCor) for BOLD and perfusion based fMRI. Neuroimage. 2007;37(1):90–101.

39. Murphy K, Fox MD. Towards a consensus regarding global signal regression for resting state functional connectivity MRI. NeuroImage. 2017;154:169–73.

40. Schaefer A, Kong R, Gordon EM, Laumann TO, Zuo XN, Holmes AJ, et al. Local-Global Parcellation of the Human Cerebral Cortex from Intrinsic Functional Connectivity MRI. Cereb Cortex. 2018;28(9):3095–114.

41. Tian Y, Margulies DS, Breakspear M, Zalesky A. Topographic organization of the human subcortex unveiled with functional connectivity gradients. Nat Neurosci. 2020;23(11):1421–32.

42. Yeo BT, Krienen FM, Sepulcre J, Sabuncu MR, Lashkari D, Hollinshead M, et al. The organization of the human cerebral cortex estimated by intrinsic functional connectivity. J Neurophysiol. 2011;106(3):1125–65.

43. Timmermann C, Roseman L, Haridas S, Rosas FE, Luan L, Kettner H, et al. Human brain effects of DMT assessed via EEG-fMRI. Proceedings of the National Academy of Sciences. 2023;120(13):e2218949120.

44. Avram M, Fortea L, Wollner L, Coenen R, Korda A, Rogg H, et al. Large-scale brain connectivity changes following the administration of lysergic acid diethylamide, d-amphetamine, and 3,4-methylenedioxyamphetamine. Molecular Psychiatry. 2024.

45. Lindquist MA, Xu Y, Nebel MB, Caffo BS. Evaluating dynamic bivariate correlations in resting-state fMRI: a comparison study and a new approach. NeuroImage. 2014;101:531–46.

46. Choe AS, Nebel MB, Barber AD, Cohen JR, Xu Y, Pekar JJ, et al. Comparing test-retest reliability of dynamic functional connectivity methods. Neuroimage. 2017;158:155–75.

47. Doss MK, Považan M, Rosenberg MD, Sepeda ND, Davis AK, Finan PH, et al. Psilocybin therapy increases cognitive and neural flexibility in patients with major depressive disorder. Translational Psychiatry. 2021;11(1):574.

48. Doss MK, May DG, Johnson MW, Clifton JM, Hedrick SL, Prisinzano TE, et al. The Acute Effects of the Atypical Dissociative Hallucinogen Salvinorin A on Functional Connectivity in the Human Brain. Scientific Reports. 2020;10(1):16392.

49. Barrett FS, Doss MK, Sepeda ND, Pekar JJ, Griffiths RR. Emotions and brain function are altered up to one month after a single high dose of psilocybin. Scientific Reports. 2020;10(1):2214.

50. Delgado-Bonal A, Marshak A. Approximate entropy and sample entropy: A comprehensive tutorial. Entropy. 2019;21(6):541.

51. Van de Bittner GC, Ricq EL, Hooker JM. A philosophy for CNS radiotracer design. Acc Chem Res. 2014;47(10):3127–34.

52. Hansen JY, Shafiei G, Markello RD, Smart K, Cox SML, Nørgaard M, et al. Mapping neurotransmitter systems to the structural and functional organization of the human neocortex. Nature Neuroscience. 2022;25(11):1569–81.

53. Wagner HH, Dray S. Generating spatially constrained null models for irregularly spaced data using Moran spectral randomization methods. Methods in Ecology and Evolution. 2015;6(10):1169–78.

54. Markello RD, Misic B. Comparing spatial null models for brain maps. NeuroImage. 2021;236:118052.

55. Bazinet V, Hansen JY, Misic B. Towards a biologically annotated brain connectome. Nature Reviews Neuroscience. 2023;24(12):747–60.

56. McIntosh AR, Lobaugh NJ. Partial least squares analysis of neuroimaging data: applications and advances. Neuroimage. 2004;23 Suppl 1:S250–63.

57. Zalesky A, Fornito A, Bullmore ET. Network-based statistic: identifying differences in brain networks. Neuroimage. 2010;53(4):1197–207.

58. Margulies DS, Ghosh SS, Goulas A, Falkiewicz M, Huntenburg JM, Langs G, et al. Situating the default-mode network along a principal gradient of macroscale cortical organization. Proceedings of the National Academy of Sciences. 2016;113(44):12574–9.

59. Sydnor VJ, Larsen B, Bassett DS, Alexander-Bloch A, Fair DA, Liston C, et al. Neurodevelopment of the association cortices: Patterns, mechanisms, and implications for psychopathology. Neuron. 2021;109(18):2820–46.

60. Carhart-Harris RL, Muthukumaraswamy S, Roseman L, Kaelen M, Droog W, Murphy K, et al. Neural correlates of the LSD experience revealed by multimodal neuroimaging. Proc Natl Acad Sci U S A. 2016;113(17):4853–8.

61. Madsen MK, Fisher PM, Burmester D, Dyssegaard A, Stenbæk DS, Kristiansen S, et al. Psychedelic effects of psilocybin correlate with serotonin 2A receptor occupancy and plasma psilocin levels. Neuropsychopharmacology. 2019;44(7):1328–34.

62. Mason NL, Kuypers KPC, Müller F, Reckweg J, Tse DHY, Toennes SW, et al. Me, myself, bye: regional alterations in glutamate and the experience of ego dissolution with psilocybin. Neuropsychopharmacology. 2020;45(12):2003–11.

63. Müller F, Holze F, Dolder P, Ley L, Vizeli P, Soltermann A, et al. MDMA-induced changes in within-network connectivity contradict the specificity of these alterations for the effects of serotonergic hallucinogens. Neuropsychopharmacology. 2021;46(3):545–53.

64. Siegel JS, Subramanian S, Perry D, Kay BP, Gordon EM, Laumann TO, et al. Psilocybin desynchronizes the human brain. Nature. 2024;632(8023):131–8.

65. Girn M, Roseman L, Bernhardt B, Smallwood J, Carhart-Harris R, Nathan Spreng R. Serotonergic psychedelic drugs LSD and psilocybin reduce the hierarchical differentiation of unimodal and transmodal cortex. Neuroimage. 2022;256:119220.

66. Roseman L, Leech R, Feilding A, Nutt DJ, Carhart-Harris RL. The effects of psilocybin and MDMA on between-network resting state functional connectivity in healthy volunteers. Front Hum Neurosci. 2014;8:204.

67. Shine JM, Lewis LD, Garrett DD, Hwang K. The impact of the human thalamus on brain-wide information processing. Nature Reviews Neuroscience. 2023;24(7):416–30.

68. Kernbach JM, Yeo BTT, Smallwood J, Margulies DS, Thiebaut de Schotten M, Walter H, et al. Subspecialization within default mode nodes characterized in 10,000 UK Biobank participants. Proceedings of the National Academy of Sciences. 2018;115(48):12295–300.

69. Schaefer A, Burmann I, Regenthal R, Arélin K, Barth C, Pampel A, et al. Serotonergic modulation of intrinsic functional connectivity. Current Biology. 2014;24(19):2314–8.

70. Tagliazucchi E, Roseman L, Kaelen M, Orban C, Muthukumaraswamy SD, Murphy K, et al. Increased Global Functional Connectivity Correlates with LSD-Induced Ego Dissolution. Curr Biol. 2016;26(8):1043–50.

71. Preller KH, Duerler P, Burt JB, Ji JL, Adkinson B, Stämpfli P, et al. Psilocybin Induces Time-Dependent Changes in Global Functional Connectivity. Biol Psychiatry. 2020;88(2):197–207.

72. Preller KH, Burt JB, Ji JL, Schleifer CH, Adkinson BD, Stämpfli P, et al. Changes in global and thalamic brain connectivity in LSD-induced altered states of consciousness are attributable to the 5-HT2A receptor. Elife. 2018;7.

73. Lord L-D, Expert P, Atasoy S, Roseman L, Rapuano K, Lambiotte R, et al. Dynamical exploration of the repertoire of brain networks at rest is modulated by psilocybin. NeuroImage. 2019;199:127–42.

74. Mortaheb S, Fort LD, Mason NL, Mallaroni P, Ramaekers JG, Demertzi A. Dynamic Functional Hyperconnectivity After Psilocybin Intake Is Primarily Associated With Oceanic Boundlessness. Biological Psychiatry: Cognitive Neuroscience and Neuroimaging. 2024;9(7):681–92.

75. Carhart-Harris RL, Friston KJ. REBUS and the Anarchic Brain: Toward a Unified Model of the Brain Action of Psychedelics. Pharmacol Rev. 2019;71(3):316–44.

76. Vollenweider FX, Preller KH. Psychedelic drugs: neurobiology and potential for treatment of psychiatric disorders. Nat Rev Neurosci. 2020;21(11):611–24.

77. Mediano PAM, Ikkala A, Kievit RA, Jagannathan SR, Varley TF, Stamatakis EA, et al. Fluctuations in Neural Complexity During Wakefulness Relate To Conscious Level and Cognition. bioRxiv. 2021:2021.09.23.461002.

78. Ort A, Smallridge JW, Sarasso S, Casarotto S, von Rotz R, Casanova A, et al. TMS-EEG and resting-state EEG applied to altered states of consciousness: oscillations, complexity, and phenomenology. iScience. 2023;26(5).

79. Mediano PAM, Rosas FE, Timmermann C, Roseman L, Nutt DJ, Feilding A, et al. Effects of External Stimulation on Psychedelic State Neurodynamics. ACS Chemical Neuroscience. 2024;15(3):462–71.

80. Leiser SC, Li Y, Pehrson AL, Dale E, Smagin G, Sanchez C. Serotonergic Regulation of Prefrontal Cortical Circuitries Involved in Cognitive Processing: A Review of Individual 5-HT Receptor Mechanisms and Concerted Effects of 5-HT Receptors Exemplified by the Multimodal Antidepressant Vortioxetine. ACS Chemical Neuroscience. 2015;6(7):970–86.

81. Juliani A, Chelu V, Graesser L, Safron A. A dual-receptor model of serotonergic psychedelics: therapeutic insights from simulated cortical dynamics. bioRxiv. 2024:2024.04.12.589282.

82. Tiger M, Varnäs K, Okubo Y, Lundberg J. The 5-HT(1B) receptor - a potential target for antidepressant treatment. Psychopharmacology (Berl). 2018;235(5):1317–34.

83. Svensson JE, Tiger M, Plavén-Sigray P, Halldin C, Schain M, Lundberg J. In vivo correlation of serotonin transporter and 1B receptor availability in the human brain: a PET study. Neuropsychopharmacology. 2022;47(10):1863–8.

84. Varnäs K, Hall H, Bonaventure P, Sedvall G. Autoradiographic mapping of 5-HT1B and 5-HT1D receptors in the post mortem human brain using [3H]GR 125743. Brain Research. 2001;915(1):47–57.

85. Montañez S, Munn JL, Owens WA, Horton RE, Daws LC. 5-HT1B receptor modulation of the serotonin transporter in vivo: studies using KO mice. Neurochemistry international. 2014;73:127–31.

86. Páleníček T, Fujáková M, Brunovský M, Horáček J, Gorman I, Balíková M, et al. Behavioral, neurochemical and pharmaco-EEG profiles of the psychedelic drug 4-bromo-2,5-dimethoxyphenethylamine (2C-B) in rats. Psychopharmacology. 2013;225(1):75–93.

87. Palamar JJ, Acosta P. A qualitative descriptive analysis of effects of psychedelic phenethylamines and tryptamines. Human Psychopharmacology: Clinical and Experimental. 2020;35(1):e2719.

88. Sayalı C, Barrett FS. The costs and benefits of psychedelics on cognition and mood. Neuron. 2023.

89. Vidaurre D, Smith SM, Woolrich MW. Brain network dynamics are hierarchically organized in time. Proceedings of the National Academy of Sciences. 2017;114(48):12827–32.

90. Menon V. Large-scale brain networks and psychopathology: a unifying triple network model. Trends Cogn Sci. 2011;15(10):483–506.

91. Valk SL, Xu T, Paquola C, Park B-y, Bethlehem RA, Vos de Wael R, et al. Genetic and phylogenetic uncoupling of structure and function in human transmodal cortex. Nature Communications. 2022;13(1):2341.

92. Mallaroni P, Mason NL, Kloft L, Reckweg JT, van Oorsouw K, Toennes SW, et al. Shared functional connectome fingerprints following ritualistic ayahuasca intake. Neuroimage. 2024;285:120480.

93. Tolle HM, Farah JC, Mallaroni P, Mason NL, Ramaekers JG, Amico E. The unique neural signature of your trip: Functional connectome fingerprints of subjective psilocybin experience. Netw Neurosci. 2024;8(1):203–25.

94. Luppi AI, Singleton SP, Hansen JY, Jamison KW, Bzdok D, Kuceyeski A, et al. Contributions of network structure, chemoarchitecture and diagnostic categories to transitions between cognitive topographies. Nature Biomedical Engineering. 2024;8(9):1142–61.

95. Escrichs A, Sanz Perl Y, Fisher PM, Martínez-Molina N, G-Guzman E, Frokjaer VG, et al. Whole-brain turbulent dynamics predict responsiveness to pharmacological treatment in major depressive disorder. Molecular Psychiatry. 2024.

96. Doss MK, Samaha J, Barrett FS, Griffiths RR, de Wit H, Gallo DA, Koen JD. Unique effects of sedatives, dissociatives, psychedelics, stimulants, and cannabinoids on episodic memory: A review and reanalysis of acute drug effects on recollection, familiarity, and metamemory. Psychological Review. 2024;131(2):523–62.

97. Ettrup A, Hansen M, Santini MA, Paine J, Gillings N, Palner M, et al. Radiosynthesis and in vivo evaluation of a series of substituted 11 C-phenethylamines as 5-HT 2A agonist PET tracers. European journal of nuclear medicine and molecular imaging. 2011;38:681–93.

98. Knudsen GM. Sustained effects of single doses of classical psychedelics in humans. Neuropsychopharmacology. 2023;48(1):145–50.

