## Supplementary Materials for "The forgotten psychedelic: Spatiotemporal mapping of brain organisation following the administration of 2C-B and psilocybin"

**Corresponding authors' emails***

**Table of Contents:**


- **S1. Methods: Study procedures**
- **S2. Methods: Subjective effect assessments**
- **S3. Methods: fMRI acquisition and preprocessing**
- **S4. Methods: Secondary entropy measures**
- **S5. Methods: Receptor mapping**
- **S6. Methods: Experiential mapping**
- **S7. Methods: Statistics**
- **S8. Methods: Cortical gradients of functional brain organisation mapping**
- **S9. Results:  Consort flowchart**
- **S10. Results:  Imaging sample demographics**
- **S11. Results:  Secondary complexity outcomes**
- **S12. Results:  Measure covariance**
- **S13. Results:  Motion assessments**
- **S14. Results:  Framewise displacement control**
- **S15. Results: Parcellation replication**
- **S16. Results: Global signal regression**
- **S17. References**

### S1. Methods: Study procedures

*Inclusion/exclusion criterion*

The inclusion criteria were: 18-40 years of age; previous experience with a psychedelic drug but not within the past 3 months; body mass index between 18 and 28 kg/m2; free from medication (any drug prescribed for a medical indication); good physical health, including the absence of major medical, endocrine, and neurological conditions; and written informed consent. Psychedelic experienced participants were recruited as to further to minimise the likelihood of serious adverse psychological effects. The exclusion criteria included history of drug abuse or addiction, pregnancy, or lactation, current or history of psychiatric disorders, absence of reliable contraceptives, previous experience of serious side effects to psychedelics, and MRI contraindications. Before inclusion, participants were screened and examined by an independent study physician, who checked for general health, conducted a resting ECG, and took blood and urine samples in which haematology, clinical chemistry and urine analyses were performed. All participants were fully informed of all procedures, possible adverse reactions, legal rights, and responsibilities, expected benefits, and their right to voluntary termination without consequences. All participants provided their informed consent, in writing, before their inclusion in the study and were remunerated for their participation.

*Screening*

Preliminary eligibility was assessed through an online pre-screening (using Qualtrics XM), which evaluated participants' prior substance use history, frequency, and geographic location. If individuals met the initial criteria, they were invited to an online briefing about the study procedures and assessments. After providing written informed consent, participants completed structured inventories that evaluated their psychiatric history (using DSM Axis I/WHO ICD-10 diagnostic classifications) and drug use history. Those who met the inclusion criteria were invited to an in-depth, in-person assessment. During this session, the study psychiatrist conducted a thorough medical screening interview to cross-check the provided information. The evaluation included a detailed review of the participant’s physical and mental health, measurement of vital signs, weight, and electrocardiogram, and a comprehensive physical exam. Laboratory tests were also conducted, including urine toxicology for illicit drugs, urinalysis, serum chemistry, haematology, and liver function tests. Participants were informed that, although they might not benefit directly from participation, their involvement could contribute to knowledge that might benefit others in the future.

*Testing Procedures*

Each experimental session lasted approximately 7 hours. On the morning of dosing, participants were instructed to eat a light breakfast at home. They were reminded to abstain from drug use, including psychedelics (for at least 3 months), alcohol (for at least 24 hours), and all other drugs of abuse (for at least 7 days before the start of testing cycles) before each experimental session. Participants were also asked to avoid caffeine and nicotine on test days. The use of psychoactive substances was prohibited for the entire 7-week duration of the study. Upon arrival on test days, the absence of drug and alcohol use was confirmed through a urine drug test and breathalyser, and a pregnancy test was administered to female participants. If all tests were negative, participants were allowed to proceed with the dosing. During the testing period, participants were permitted to interact with the investigator, rest, read, or listen to music via headphones. They remained under supervision until the session ended and the experimenters determined they were fit to leave.

### S2. Methods: Subjective effect assessments

*Subjective effect visual analogue scales (VAS)*

Subjective effects were assessed repeatedly using visual analogue scales (VASs) at baseline (0h), +0.5, +1, +1.5, +2, +3, +4, +5, +6 hours after drug administration. These scales included the primary VAS items: "*any drug effect*, and "*drug high*", presented as 100-mm horizontal lines (0–100%), marked from "*not at all" on* the left to "*extremely*" on the right (1). The VAS "*any drug effect*" is an overall effect measure to characterise the overall effect intensity and time course. Prior dose-effect studies have previously demonstrated it to useful marker for pharmacokinetic-pharmacodynamic modeling of psilocybin and LSD's effects (2, 3). Separately, "any drug effect" is interrelated with "drug high", a measure of stimulating effects. These items were used to define dosage equivalence from previous data (4, 5) and found to yield psychotropic equivalence in the complete behavioural sample across a range of measurements (6, 7).

*Altered States of Consciousness Rating Scale (5D-ASC)*

The 5D-ASC (8) consists of 94 retrospective 100-mm VAS. The ASC items are grouped into five main dimensions comprising 11 lower-order scales (1) 'oceanic boundlessness' (OB) measures derealisation and depersonalisation accompanied by changes in affect ranging from heightened mood to euphoria and/or exaltation as well as alterations in the sense of time. The corresponding subfactors include "experience of unity," "spiritual experience," "blissful state," "insightfulness," and "disembodiment." (2) 'anxious ego dissolution' (AED) measures ego disintegration associated with loss of self-control, thought disorder, arousal, and anxiety. The corresponding subfactors comprise "impaired control of cognition" and "anxiety." (3) 'visionary restructuralisation' (VR) refers to 'elementary hallucinations', 'visual (pseudo-) hallucinations', 'synaesthesia', 'changed meaning of percepts', 'facilitated recollection', and 'facilitated imagination'. It consists of the lower-order scales "complex imagery," "elementary imagery," "audio-visual synesthesia," and "changed meaning of percepts." (4) 'auditory alterations' (AA) refers to acoustic hallucinations and distortions in auditory experiences and (5) the dimension 'reduction of vigilance' (RV) relates to states of drowsiness, reduced alertness, and related impairment of cognitive function. The 5D-ASC is the most widely used psychometric assessment of classical psychedelic and entactogenic effects and has been deployed across a range of altered states of consciousness (9).

*The Ego-Dissolution Inventory (EDI)*

The Ego-Dissolution Inventory is a self-reported scale used to specifically assess subjective feelings of ego-dissolution/loss of self after drug intake (10). The questionnaire consists of 8 VAS items (100-mm) which participants have to rate retrospectively, including: I experienced a dissolution of my "self" or ego; I felt at one with the universe; I felt a sense of union with others; I experienced a decrease in my sense of self-importance; I experienced a disintegration of my "self" or ego; I felt far less absorbed by my own issues and concerns; I lost all sense of ego; All notion of self and identity dissolved away. The EDI has been shown to relate to the underlying effects of 5-$HT_{2A}$ agonists on functional brain organisation (11, 12).

*Amsterdam Resting-State Questionnaire (ARSQ)*

Following scanner exit, participants were asked to complete the ARSQ to gauge potential between-condition differences in the degree of arousal, mind wandering and thought content during resting state acquisition. The ARSQ is a 50-item list of Likert-type statements rated on a scale from 1 ("Completely disagree") to 5 ("Completely agree") probing potential subjective experiences arising during the resting stating and clusters into 7 dimensions; Discontinuity of Mind, Theory of Mind, Self, Planning, Sleepiness, Comfort, and Somatic (13). The ARSQ has been validated during EEG and fMRI resting-state with the primary mentation subscales; Discontinuity of Mind, Theory of Mind, Self, and Planning being shown to be stable across different experimental settings (13-15,16)

### S3. Methods: fMRI acquisition and preprocessing

A 32 receiving-channel head array Nova coil (NOVA Medical Inc., Wilmington MA) was used to acquire imaging data for all participants on the 7T MAGNETOM scanner for an imaging session which spanned a time range of ± 75 to ± 130 min after administration. Scanning comprised structural imaging, magnetic resonance spectroscopy, task and rsfMRI.

Structural T1-weighted (T1w) images were collected at the beginning of each imaging session at an isotropic resolution of 0.9mm using a magnetisation-prepared 2 rapid acquisition gradient-echo (MP2RAGE) acquisition (190 sagittal slices following parameters: repetition time (TR) = 4,500 ms, echo time (TE) = 2.39 ms, inversion times TI1/TI2 = 900/2750 ms, flip angle1 = 5°, flip angle2 = 3°, voxel size = 0.9 mm isotropic, bandwidth = 250 Hz/pixel). High-resolution blood oxygenation level-dependent (BOLD) signal estimation was performed using an echo planar imaging sequence of 1.5 mm isotropic voxel size (TR = 1400 ms; TE = 21 ms; field of view=198 mm; flip angle = 60°; oblique acquisition orientation; interleaved slice acquisition; 72 slices; slice thickness = 1.5 mm).

Resting-state data (516 volumes, 12 minutes) were acquired approximately 110 minutes after substance intake and were immediately followed by five EPI volumes in the opposite phase direction for magnetic susceptibility distortion correction. During this time, participants were shown a black fixation cross on a white background and were instructed *to "Please look at the cross at all times and try to clear your mind. Stay as still as possible and do not fall asleep"* as per prior work (12, 17).

*Anatomical data preprocessing*
Across all anatomical scans found for a given subject (across all conditions), the following preprocessing was performed. 7T MP2RAGE signal inhomogeneity was normalised by reconstructing "robust" T1w equivalents. In sum, as per (17) a normalised complexity ratio was extrapolated from T1w (GRETI1) and

PDw (GRETI2) image volumes and applied to generate a uniform T1w image volume of minimal signal intensity variance. All images were then corrected for intensity non-uniformity (INU) with N4BiasFieldCorrection, distributed with ANTs 2.3.3. The T1w-reference was then skull-stripped with a Nipype implementation of the antsBrainExtraction.sh workflow (from ANTs), using OASIS30ANTs as target template. Brain tissue segmentation of cerebrospinal fluid (CSF), white-matter (WM) and gray-matter (GM) was performed on the brain-extracted T1w using fast (FSL 6.0.5.1:57b01774, RRID:SCR_002823]). A T1w-reference map was computed after registration of 3 T1w images (after INU-correction) using mri_robust_template (FreeSurfer 7.2.0). Brain surfaces were reconstructed using recon-all (FreeSurfer 7.2.0, RRID:SCR_001847]), and the brain mask estimated previously was refined with a custom variation of the method to reconcile ANTs-derived and FreeSurfer-derived segmentations of the cortical gray-matter of Mindboggle (RRID:SCR_002438). Volume-based spatial normalization to two standard spaces (MNI152NLin2009cAsym, MNI152NLin6Asym) was performed through nonlinear registration with antsRegistration (ANTs 2.3.3), using brain-extracted versions of both T1w reference and the T1w template. The following templates were selected for spatial normalization: ICBM 152 Nonlinear Asymmetrical template version 2009c [RRID:SCR_008796; TemplateFlow ID: MNI152NLin2009cAsym], FSL's MNI ICBM 152 non-linear 6th Generation Asymmetric Average Brain Stereotaxic Registration Model [ RRID:SCR_002823; TemplateFlow ID: MNI152NLin6Asym].

*Functional data preprocessing*

Across all scans for a given subject (across all conditions), the following functional preprocessing was performed. The initial two volumes were excluded, and a B0-nonuniformity map (fieldmap) was estimated based on the two echo-planar imaging (EPI) references with topup (FSL 6.0.5.1:57b01774) to correct for non-steady state and magnetic susceptibility distortion. First, a reference volume and its skull-stripped version were generated using a custom methodology of fMRIPrep. Head-motion parameters with respect to the BOLD reference (transformation matrices, and six corresponding rotation and translation parameters) are estimated before any spatiotemporal filtering using mcflirt (FSL 6.0.5.1:57b01774). The estimated fieldmap was then aligned with rigid registration to the target EPI (echo-planar imaging) reference run. The field coefficients were mapped on to the reference EPI using the transform. BOLD runs were slice-time corrected using 3dTshift from AFNI ( RRID:SCR_005927). The BOLD reference was then co-registered to the T1w reference using bbregister (FreeSurfer) which implements boundary-based registration. Co-registration was configured with six degrees of freedom. All resamplings can be performed with a single interpolation step by composing all the pertinent transformations (i.e. head-motion transform matrices, susceptibility distortion correction when available, and co-registrations to anatomical and output spaces). Gridded (volumetric) resamplings were performed using antsApplyTransforms (ANTs), configured with Lanczos interpolation to minimize the smoothing effects of other kernels. Non-gridded (surface) resamplings were performed using mri_vol2surf (FreeSurfer).

### S4. Methods: Secondary entropy measures

Following the recommendations of McCulloch et al., we calculated additional complexity measures according to their a) scale of organisation b) level of reproducibility and c) their association with psychedelic effects. Note the degree of associative validity of these measures in the aforementioned work is limited by the use of an ascending dose design without the use of a placebo control.

*Edge-wise Shannon entropy of dynamic conditional correlations (dccEn)*

Following the generation of DCC timeseries, the probability distribution for each ROI-to-ROI DCC time series was determined, and Shannon entropy was calculated across bins using MATLAB's histcounts function. The Shannon entropy for each ROI pair (i, j) is:

$$H(i, ii) = -\sum_{k} P_k log P_k$$

Where $P_k$ are the probabilities obtained from histcounts, representing the likelihood of observing each value of the DCC between ROI *i* and ROI j. The $\Sigma_k$ is over all bins (or distinct DCC values) in the probability histogram. dccEn measures the uncertainty in the dynamic connectivity between each pair of ROIs over time. Higher entropy values indicate more diversity in the DCC matrix (less predictability), while lower entropy values suggest more consistent, predictable connectivity.

*Whole-brain path-length degree distribution (degreeEn)*

Degree refers to the count of non-zero elements in any row of a thresholded matrix. Static functional connectivity matrices were thresholded based on mean-degree criteria, using absolute correlation values, without applying a p-value threshold. The thresholding process involved two steps. Initially, any correlation with a corresponding p-value greater than 0.05 was set to zero. The next step aimed to achieve a predetermined mean degree across rows. To accomplish this, the threshold was gradually increased until the mean number of non-zero elements reached the desired level. A scan-specific threshold was applied to achieve a mean degree of 27, as this threshold produced the most significant effect in the original study where it was used (18).

Consequently, each scan may have a different absolute threshold value, but the mean degree remains consistent. The final entropy calculation involved determining the Shannon entropy of the degree distribution across ROIs. The matrix was then binarised by setting all non-zero elements to one. Finally, the 'shortest path length,' defined as the minimum number of edges required to connect any two nodes, was computed, followed by calculating the Shannon entropy of the distribution of path lengths from each node to all others.

*Whole-brain Lempel Ziv complexity (zivEn)*

The BOLD time series for each ROI were first transformed using the Hilbert method. The amplitude of the resulting Hilbert series was then binarised by comparing it to the mean amplitude for that region—values greater than the mean were assigned '1,' and those less were assigned '0.' These binarised time series were combined into a T × N matrix, where N represents the number of regions and T represents the number of time points. This matrix was then collapsed into a single vector to compute the Lempel-Ziv complexity over time (using the LZ76 algorithm) by concatenating the regional time series. This calculation reflects the temporal entropy across ROIs, as detailed in (19, 20).

It is worth considering that independent of their selection choice, each complexity measure might offer a distinct approach to charting the same construct of 'brain complexity': **sampEn** reflects regional complexity through the unpredictability of each region's BOLD time series sequence; **dccEn** captures edge-based temporal complexity; **degreeEn** indicates the complexity of whole-brain connectivity motifs; and **zivEn** assesses the complexity of a whole-brain time series by evaluating the "compressibility" of its BOLD time series sequence.

### S5. Methods: Receptor mapping

Dominance analysis involves fitting the same regression model to every possible combination of predictors (creating $2^p - 1$ submodels for a model with p predictors) (21). Total dominance is characterised as the mean increase in $R^2$ when incorporating a single predictor of interest to a submodel, considering all $2^p - 1$ submodels. The aggregate dominance of all input variables equals the total adjusted $R^2$ of the comprehensive model, rendering the relative importance percentage an easily understandable technique that apportions the overall effect size among predictors. As a result, in complement to alternative methods for evaluating predictor significance, such as univariate correlations, dominance analysis takes into account predictor-predictor interactions and offers interpretability.

Moran spectral randomisation relies on a spatially informed weight matrix *W* (22). The eigenvectors of W provide an estimate of the autocorrelation in the brain and are used to impose a similar spatial structure on random, normally distributed surrogate data (10 000 permutations). Here, W is defined as the inverse of the distance matrix between brain regions. Data was generated separately for each hemisphere using the same random seed to obtain null annotations that preserve homotopy across hemispheres.

### S6. Methods: Experiential mapping

PLS analyses are best suited for instances where the number of predictors exceeds the number of observations and when the predictors (regional coherence scores) exhibit multicollinearity (23). Using a singular value decomposition, behavioural PLS attempts to find linear combinations of features from the provided arrays that maximally covary with each other. The decomposition is performed on the cross-covariance matrix $R$, where $R= YT \times X$ which represents the covariation of all the input features across samples. To assess the significance of latent variable (LV) multivariate patterns, we computed permutation tests (10,000 permutations). Each permutation shuffled the order of the observations of the subjective effect matrix before running PLS on the resulting surrogate data under the null hypothesis that there was no relationship between the subjective effect data and neural data. A *p*-value for each LV was computed as the proportion of times the permuted singular values exceeded that of the original data. We also estimated the stability of regional coherence loadings by bootstrapping observations with replacement (10,000 bootstraps). Generally, overall model in-sample correlation scores can be inflated in the absence of external validation approaches, which seek to compare model performance using "unseen" or "out-of-sample" data. Here, limited by the small number of observations - we attempted to quantify overall model bias by performing a *k*-fold cross-validation approach across observations. Specifically, we used 5-fold cross-validation by setting test splits to 5, where the dataset was randomly partitioned into five subsets. In each fold, 75% of the data was used for training and 25% for testing, ensuring robust evaluation across multiple splits. The training data and corresponding group assignments (2C-B, psilocybin) were used to perform singular value decomposition (SVD). SVD decomposed the training data into its component matrices, revealing the latent structure between the input variables (X) and the output variables (Y). After the initial decomposition, test set data is based on the structure learned from the training set. Rescaling is performed for each group, concatenating the final test predictions into a single matrix. From these predictions, we derived out-of-sample correlations. This measured the correlation between the predicted values and the actual test set values, indicating how well the model's predictions aligned with the true data.

### S7. Methods: Cortical gradients of functional brain organisation mapping

Current theories suggest the transmodal axis of hierarchical brain organisation to be important for psychedelic effects (24, 25). To examine whether this extends to neurobehavioural markers of subjective effects, we assessed the spatial concordance of our overall model PLS bootstrap ratios and the primary gradient of FC identified by Margulies et al (26) and the Sensorimotor-Association (S-A) axis by Sydnor et al. (36) using spin-permuted (10 000 permutations) Pearson correlations.

The diffusion-embedding decomposition of human functional connectivity (FC) reveals a principal gradient, which spans between two extremes: unimodal regions responsible for primary sensory and motor functions, and transmodal regions, known in humans as the default-mode network (DMN). These DMN regions are situated at the maximal geodesic distance along the cortical surface, symmetrically positioned relative to primary sensory and motor areas. The principal gradient offers a spatial organisational framework that underpins multiple large-scale networks and delineates a continuum from unimodal to hetero-modal activity, as evidenced by functional meta-analyses (26). The principal gradient was obtained from neuromaps (https://github.com/netneurolab/neuromaps/) and parcellated into 200 regions of the surface-based Schaefer Yeo-17 atlas (27, 28).

The Sensorimotor-Association (S-A) axis was developed by averaging the rank orderings of ten cortical feature maps, each reflecting systematic variations across lower-order sensorimotor, middle-order unimodal and multimodal, and higher-order heteromodal and paralimbic association cortices (36). These feature maps include: the functional hierarchy identified by the principal gradient of functional connectivity, the evolutionary hierarchy shown through cortical areal expansion from macaques to humans, the anatomical hierarchy measured by the adult T1w/T2w ratio, local areal scaling related to brain size, aerobic glycolysis representing brain metabolism, cerebral blood flow reflecting perfusion, gene expression indexed by the principal component of brain-expressed genes, a functional brain pattern captured by NeuroSynth meta-analytic decodings, a histological gradient of cytoarchitecture derived from the BigBrain atlas, and cortical thickness measured via structural MRI. This combined S-A axis highlights a major motif of cortical organization, reflecting the progression from primary sensorimotor areas (lower ranks) to higher-order association regions in the frontal, temporal, and parietal lobes (higher ranks). The S-A axis was obtained from neuromaps (https://github.com/netneurolab/neuromaps/) and parcellated into 200 regions of the surface-based Schaefer Yeo-17 atlas (27, 28).

These vectors were correlated with our latent component regional bootstrap ratios, and FDR-corrected significance tested against spin-permutation null models. Spin permutation testing allows for the mitigation of potential confounding effects of spatial autocorrelations when assessing relationships between two cortical maps (29). Spatial maps were subject to 10,000 random spherical rotations at a parcel level to generate null models of spatial alignment. Permutation *p* values ($p_{FDRspin}$) values were computed as the proportion of null values of the intermodal Pearson correlation coefficient greater than the real values of the correlation coefficient. For the corresponding toolbox, see https://github.com/frantisekvasa/rotate_parcellation.

### S8. Methods: Statistics

Network-based statistics (NBS) is a nonparametric statistical method is designed to control the family-wise error due to multiple comparisons, specifically for application to graph data (30). Connected components of the graph are identified from edges that survive a statistical threshold (F-test; here calculated apriori according to the study's degrees of freedom [1,58] $f$ = 4.09). In turn, the statistical significance of such connected components is estimated by comparing their topological extension against a null distribution of the size of connected components obtained from non-parametric permutation testing (10 000 permutations). This approach rejects the null hypothesis on a component basis and therefore achieves superior power compared to mass-univariate approaches.

### S9. Results: CONSORT flow chart

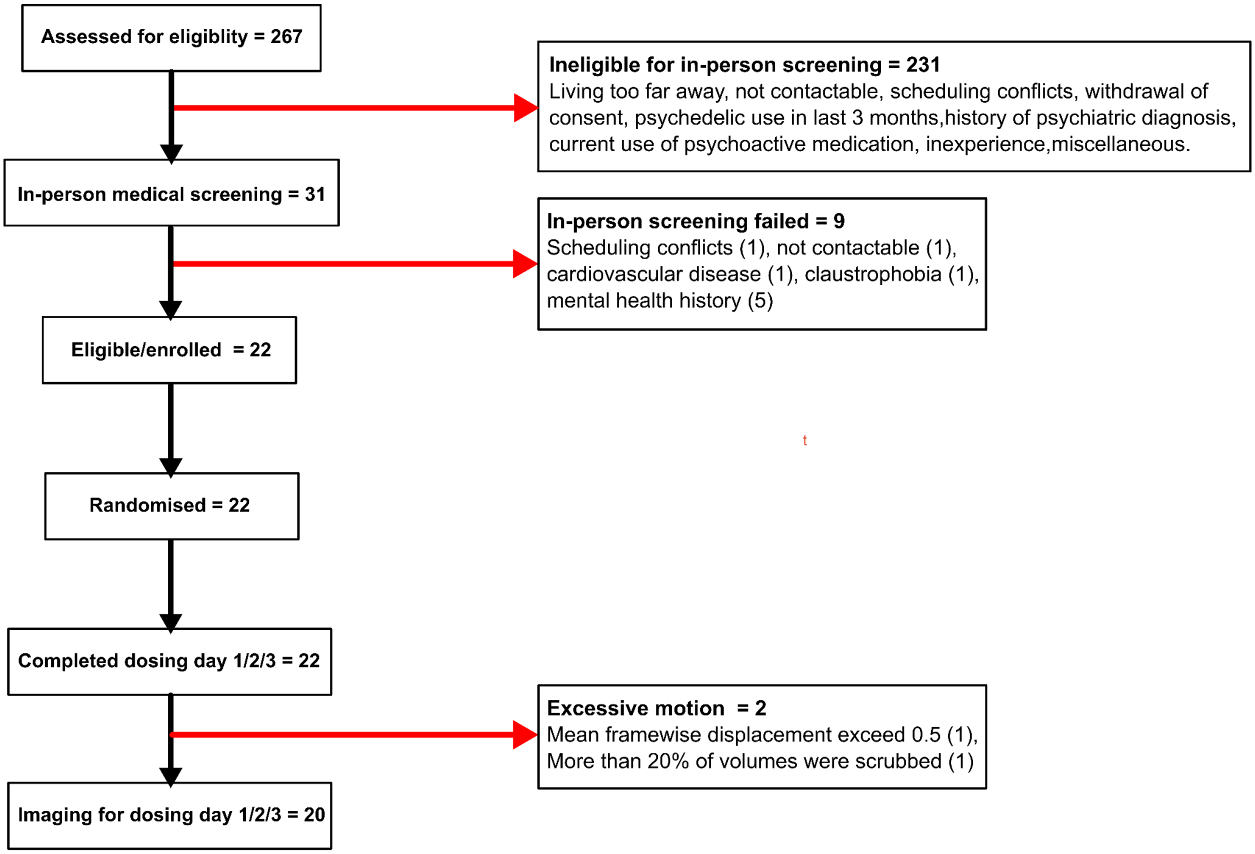


**Figure S1. Study inclusion schematic.** Adapted from original study publication (6)

### S10. Results: Imaging sample demographics

For the complete (N = 22) study demographics, see the corresponding article (6).

| **Demographics (N = 20)** | ***Mean (SD)*** |
|---|---|
| Sex (male/female), n, total | 10/10 |
| Age, years | 25.2 (4.42) |
| Weight, kg | 69.49 (10.47) |
| History of psychedelic use, years | 3.78 (1.78) |
| Lifetime psychedelic use, number of occasions | 26.16 (42.53) |
| Cannabis consumption, per month | 2.28 (2.03) |
| Alcohol consumption, glasses per week | 2.92 (3.86) |
| Caffeine consumption, units per week | 12.76 (10.12) |
| Nicotine consumption, cigarettes per week | 3.35 (7.34) |

### S11. Results: Secondary complexity outcomes

We calculated three additional measures of signal complexity (dccEn, zivEn and degreeEn, see Figure S2). No main effect of drug was identified for either dccEn (NBS max edge *f*-stat: 15.94, $p_{NBS}$ >0.05) or whole-brain degreeEn ($f_{(1,58)}$ = 0.78, $p$ = 0.3801). LMEMs identified a significant effect of drug for whole-brain zivEn ($f_{(1,58)}$ = 10.50, $p$ = 0.0019) with FDR-corrected posthoc testing reflecting significantly increased values for each drug relative to placebo (2C-B > Placebo: *t*-stat = 3.21, $p_{FDR}$ = 0.0138, Psilocybin > Placebo: *t*-stat = 2.75, $p_{FDR}$ = 0.0193) and no significant differences between conditions (Psilocybin > 2C-B: *t*-stat = -0.29 $p_{FDR}$ = 0.7721 ).

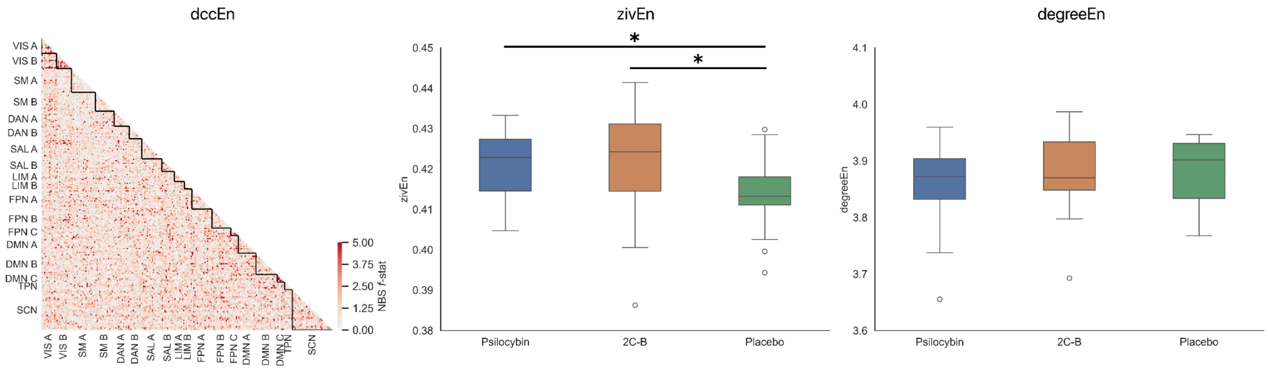


**Figure S2. Secondary outcome measures of brain complexity**. For each edge, *f*-statistics corresponding to the effect of drug (non-significant $p$ > 0.05) following the assessment of dccEn are plotted in the matrix. Cortical networks are based on the 17-network, 200-parcel Schaefer-Yeo parcellation (Schaefer et al. 2017) augmented with 32 subcortical regions from Tian et al. 2020. Red indicates the strength of the effect of drug in NBS. Regions are ordered left and then right within each network. Boxplots represent whole brain values for zivEn and degreeEn with asterixes corresponding to FDR-corrected posthoc *t*-tests following a main effect of drug. Significance is denoted as follows: * $p$ < 0.05, ** $p$< 0.01, *** $p$ < 0. 001.

### S12. Results: Measure covariance

To determine if changes in primary measures of functional organisation (sFC, dFC, sampEn) used for modelling are not necessarily linearly and/or highly correlated we a) performed FDR-corrected Pearson correlations across mean regional outcomes values (regions x features) and b) performed $FDR_{Moran}$-corrected Pearson correlations for each contrast's outcome raw nodal sum *t*-statistics. As shown in Figure S3, while raw measures of dFC, sFC and sampEn show small-moderate associations in a condition-dependent manner, these do not necessarily mean they follow one another when looking at drug-induced effects. It is also perhaps worth noting - based on the findings of Doss et al. (albeit with a different complexity measure), that the presented inverse associations below between eFC and dFC were context-dependent.

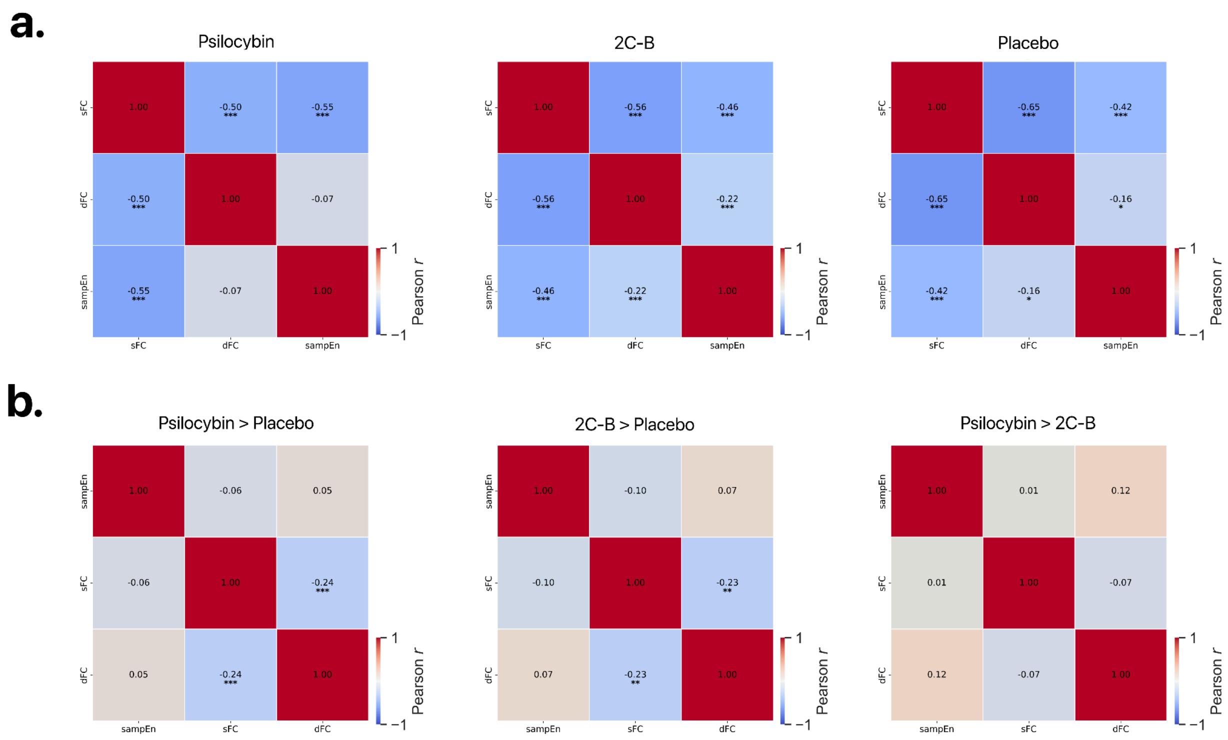


**Figure S3. Measure covariance**. A) Nodal values for each primary outcome measure were averaged per condition and their covariance was assessed using Pearson correlations (feature x regions rather than the coherence matrix regions x features). FDR correction was applied to correct for multiple comparisons. B) Nodal *t*-statistics for each equivalent measure per statistical contrast were subjected to an equivalent approach (regions x feature *t*-statistic). Moran's spectral randomisation (10 000 permutations) was employed to derive significance values prior to FDR correction. Significance is denoted as follows: * $p_{FDR} < 0.05$, ** $p_{FDR} < 0.01$, *** $p_{FDR} < 0.001$.

### S13. Results: Motion assessments

Following LMEMs, neither scrubbing nor mean framewise displacement (FD) exhibited a significant effect of drug (Scrub:$f_{(1,58)}$ = 0.76, $p$ = 0.4700; FD: $f_{(1,58)}$ = 1.41, $p$ = 0.2532). We also derived Pearson correlations per condition for each subject's regional (sampEn, gFC) and edge measures (sFC and dFC) vs FD. These did not yield any significant associations (see density plots in Figure S4b and S4c). Note that these were corrected for multiple comparisons using FDR for the number of possible regions/edges given the extreme number of possible combinations but not across conditions or measures. Condition-dependent distance-based associations were noted for our connectomic measures (see Figure S4b)

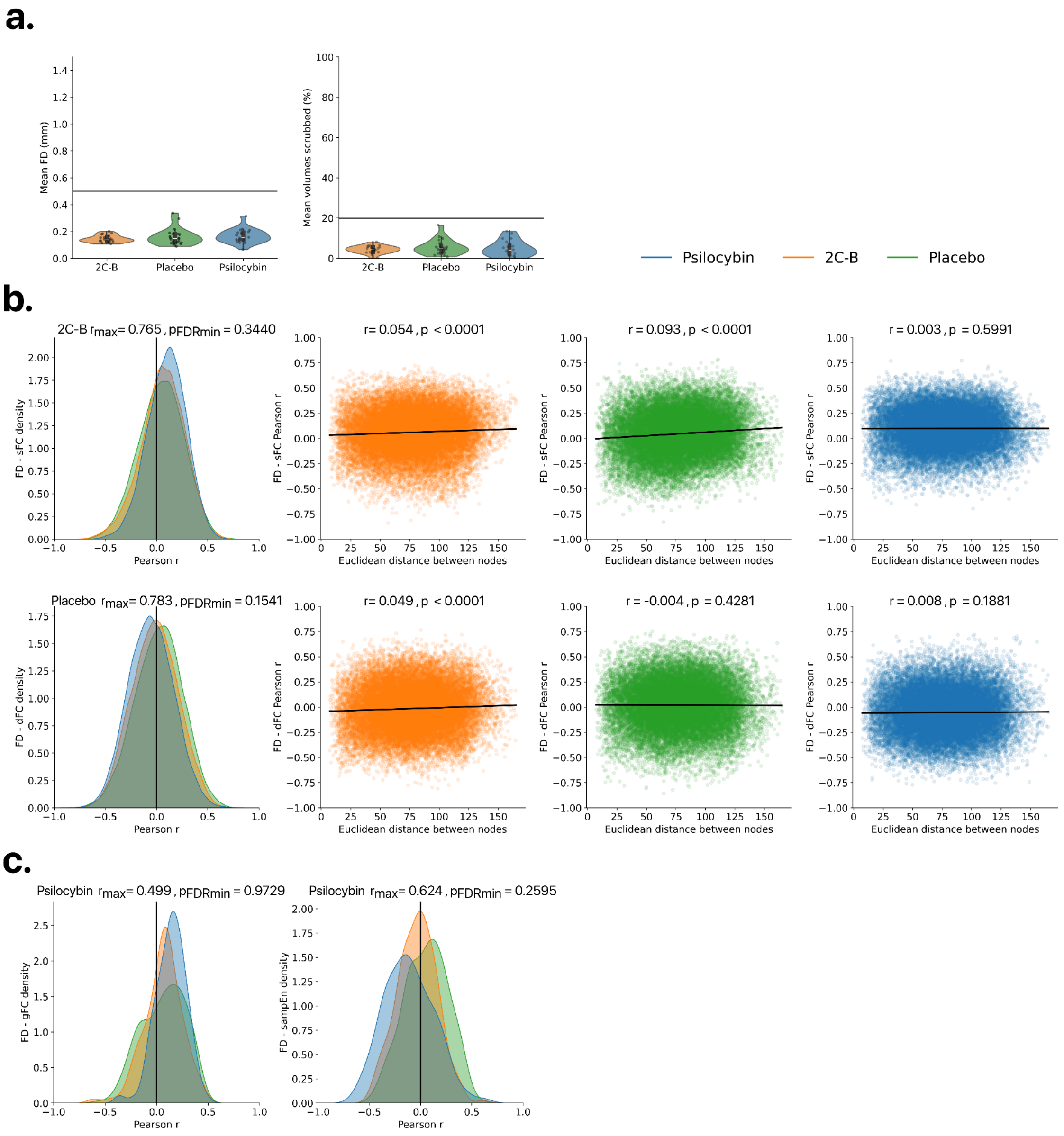

**Figure S4. Motion assessments**. A) Violin plots of scrubbing and mean framewise displacement values per condition and according to their respective thresholds. B) Connectomic quality control assessments. Kernel density plots represent the distribution of Pearson *r* values following the correlation of all unique possible fisher-transformed FC edge values per subject and corresponding framewise displacement, per session. The most significant correlations are plotted following FDR correction across all edges. Scatter plots display the relationship between Euclidian distance between brain nodes and motion vs functional connectivity. C) Regional quality control assessments. Kernel density plots represent the distribution of Pearson *r* values following the correlation of all regional values per subject and corresponding framewise displacement, per session. The most significant correlations are plotted following FDR correction across all regions.

### S14. Results: Framewise displacement control

To further assess the influence of motion, we repeated all primary analyses incorporating framewise displacement (FD) as a regressor of no interest in our second-level analyses. As per Figure S5a, connectomic measures showed good consistency following FD inclusion for sFC (231 nodes, 1927 edges, $p_{NBS}$ = 0.0465) and dFC (231 nodes, 1760 edges, $p_{NBS}$ = 0.0128), no main effect once more for dccEn ($p_{NBS}$> 0.05). Regional measures were generally topographically similar (see Figure S5b).

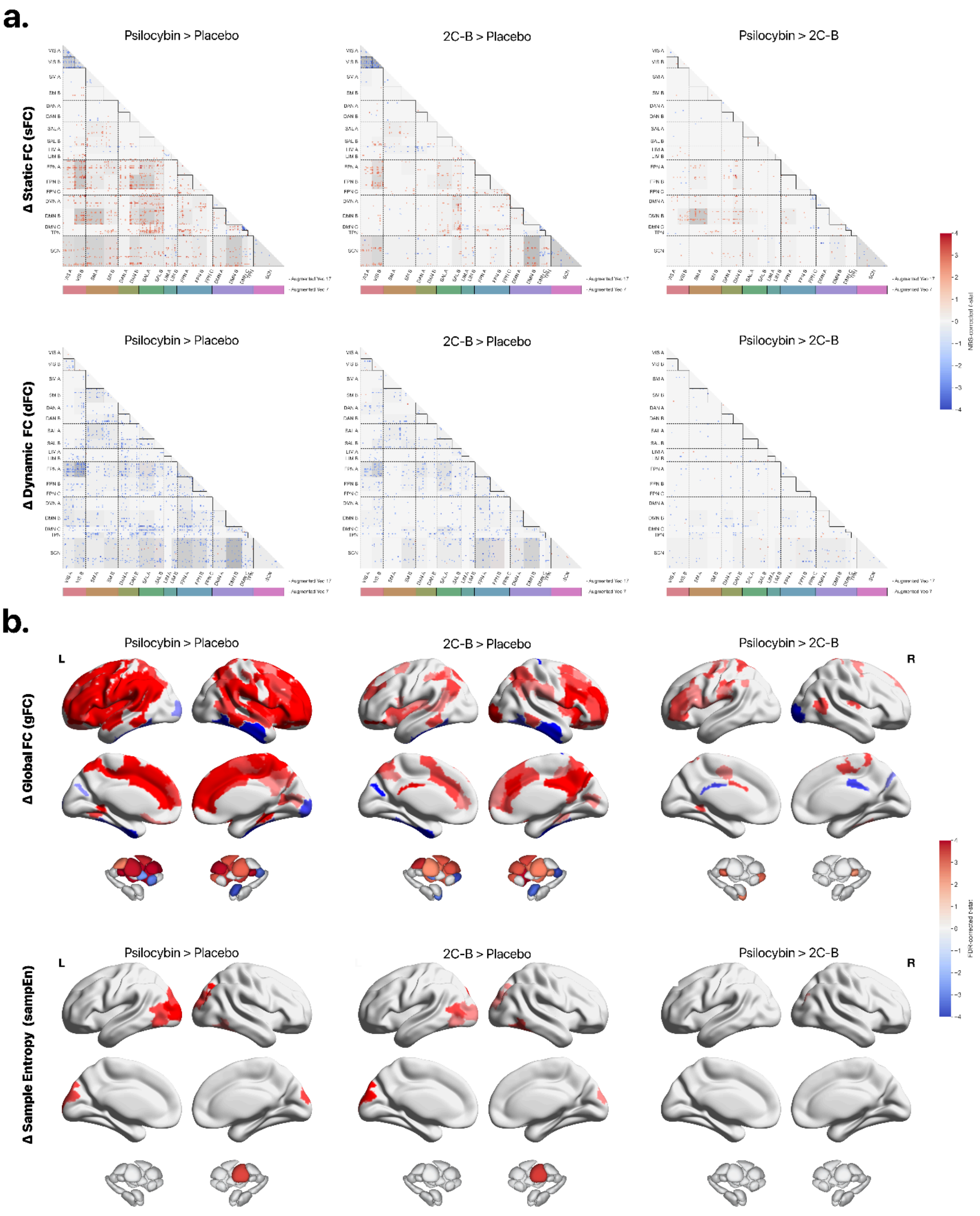

**Figure S5. rsfMRI psychedelic effect benchmarks controlling for framewise displacement**. A) Significant differences in drug-mediated static and dynamic FC across all contrasts. Cortical networks are based on the 17-network, 200-parcel Schaefer-Yeo parcellation (Schaefer et al. 2017) augmented with 32 subcortical regions from Tian et al. 2020. Red indicates drug-mediated increases in interregional FC, and blue indicates decreases. Grey hues represent the proportion of significant edges for a given Yeo 17 network pair relative to the total number of significant edges. Regions are ordered left and then right within each network. B) Significant differences in drug-mediated regional gFC (i.e., the average of a given seed region's FC to the rest of the connectome) and sampEn (the complexity of a regional timeseries) across all contrasts. Abbreviations: VIS, visual; SMN, somatomotor network; DAN, dorsal attention network; SAL, salience network; LIM, limbic; FPN, frontoparietal network; DMN, default mode network ; TPN (temporoparietal network); SCN (subcortical network).

### S15. Results: Parcellation replication

Results showed good topographical consistency at a finer-grain Schaefer 454 spatial resolution (see Figure S6). All analyses were performed identically to those used to derive our primary findings. NBS analyses revealed a significant effect of drug for sFC ( 452 nodes, 6549 edges, $p_{NBS}$ = 0.0341) as well as dFC (454 nodes, 5873 edges, $p_{NBS}$ = 0.0139) with no main effect once more for dccEn ($p_{NBS}$ > 0.05) Regional analyses showed similar results, with sparser PFC differences between Psilocybin and 2C-B.

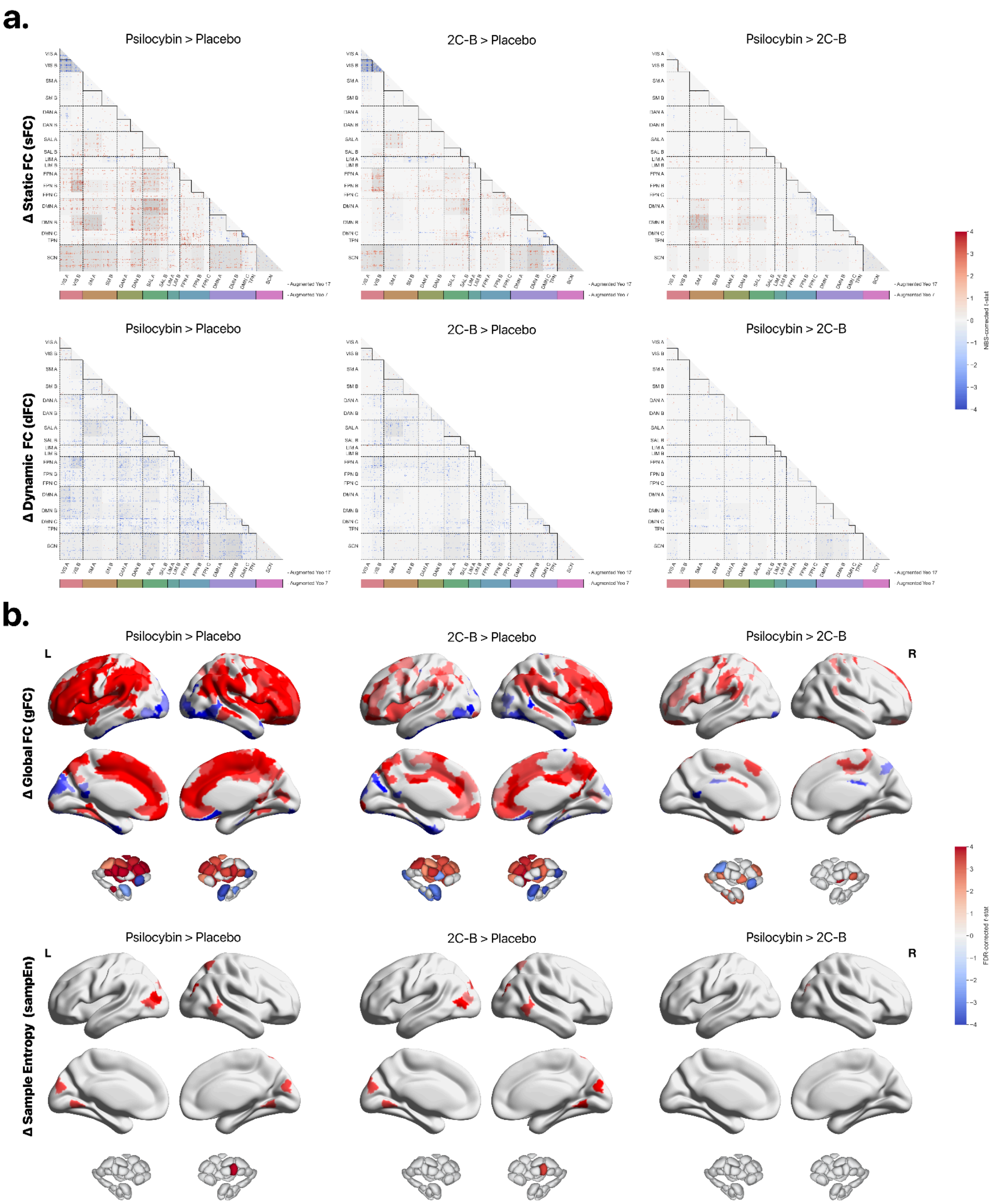

**Figure S6. rsfMRI psychedelic effect benchmarks at a finer scale**. A) Significant differences in drug-mediated static and dynamic FC across all contrasts. Cortical networks are based on the 17-network, 400-parcel Schaefer-Yeo parcellation (Schaefer et al. 2017) augmented with 54 subcortical regions from Tian et al. 2020. Red indicates drug-mediated increases in interregional FC, and blue indicates decreases. Grey hues represent the proportion of significant edges for a given Yeo 17 network pair relative to the total number of significant edges. Regions are ordered left and then right within each network. B) Significant differences in drug-mediated regional gFC (i.e., the average of a given seed region's FC to the rest of the connectome) and sampEn (the complexity of a regional timeseries) across all contrasts. Abbreviations: VIS, visual; SMN, somatomotor network; DAN, dorsal attention network; SAL, salience network; LIM, limbic; FPN, frontoparietal network; DMN, default mode network ; TPN (temporoparietal network); SCN (subcortical network).

### S16. Results: Global signal regression

Global signal regression (GSR) is a controversial preprocessing step that removes the time series of signal intensity averaged across all brain voxels through linear regression. To assess its differential influence on our results, GSR was incorporated during denoising by employing subject-level grey-matter masks atop existing nuisance regressors during aCompCor denoising. While GSR is considered to be an absolutist approach to removing possible motion artefacts, GSR can induce spurious anticorrelations and may also remove meaningful neural information (31). For example, GSR can alter the topographical distribution of transmodal sFC anticorrelations and disproportionally affect its derivative gFC according to data acquired following acute psychedelic administration (32-34).

Following its inclusion we find the most salient reported outcomes were preserved after its use, with some secondary effects (see Figure S7). NBS analyses revealed an extensive main effect of drug for sFC (232 nodes, 1691 edges, $p_{\text{NBS}} < 0.0001$) and dFC (230 nodes, 1382 edges $p_{NBS} = 0.0118$), with no significant effect for dccEn ($p_{\text{NBS}} > 0.05$). Whereas our primary findings survived its use - visually, the inclusion of GSR introduced significant sFC reductions in broader transmodal within-network ($DMN_{A/B/C}$ & $FPM_{A/B/C}$) connectivity for each drug-placebo contrast and strengthened existing findings of increased between-network connectivity. Differences between 2C-B and psilocybin were found to be more extensive following its use with network pairs such as $SMN_A$ - $SAL_A$ showing greater sFC under psilocybin. Findings for dFC were largely consistent with the absence of its use, showing extensive reductions across the whole brain. Regional sampEn findings were also preserved, showing increases in visual cortices and thalamic nuclei for each drug.

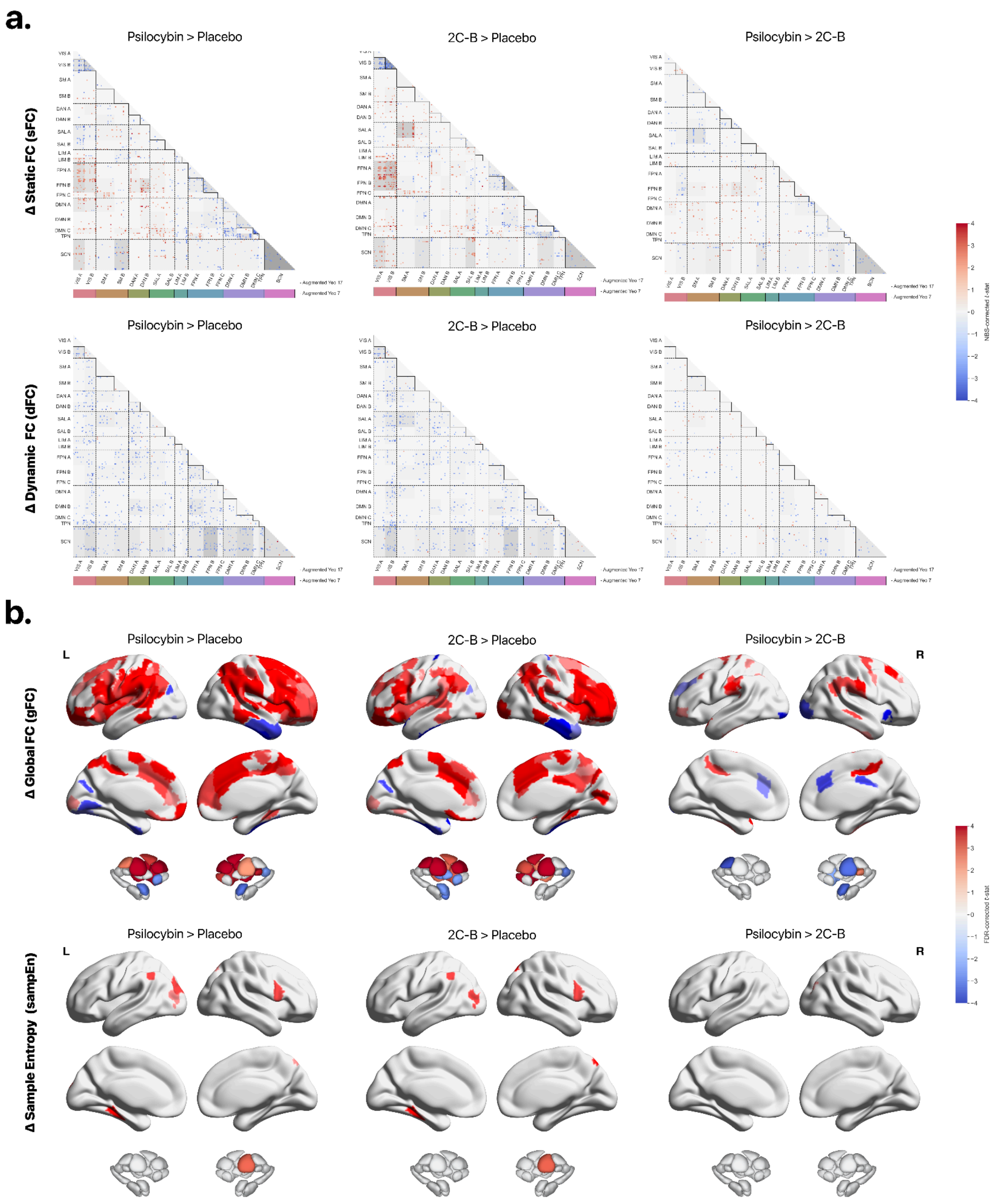


**Figure S7. rsfMRI psychedelic effect benchmarks following the use of global signal regression**. A) Significant differences in drug-mediated static and dynamic FC across all contrasts. Cortical networks are based on the 17-network, 200-parcel Schaefer-Yeo parcellation (Schaefer et al. 2017) augmented with 32 subcortical regions from Tian et al. 2020. Red indicates drug-mediated increases in interregional FC, and blue indicates decreases.

Grey hues represent the proportion of significant edges for a given Yeo 17 network pair relative to the total number of significant edges. Regions are ordered left and then right within each network. B) Significant differences in drug-mediated regional gFC (i.e., the average of a given seed region's FC to the rest of the connectome) and sampEn (the complexity of a regional timeseries) across all contrasts. Abbreviations: VIS, visual; SMN, somatomotor network; DAN, dorsal attention network; SAL, salience network; LIM, limbic; FPN, frontoparietal network; DMN, default mode network; TPN (temporoparietal network); SCN (subcortical network).

Given extensive discussions on the use of GSR to analyse global connectivity measures and its corresponding effects on static functional connectomes, we more closely assessed how it may differentially affect each compound (see Figure S7).

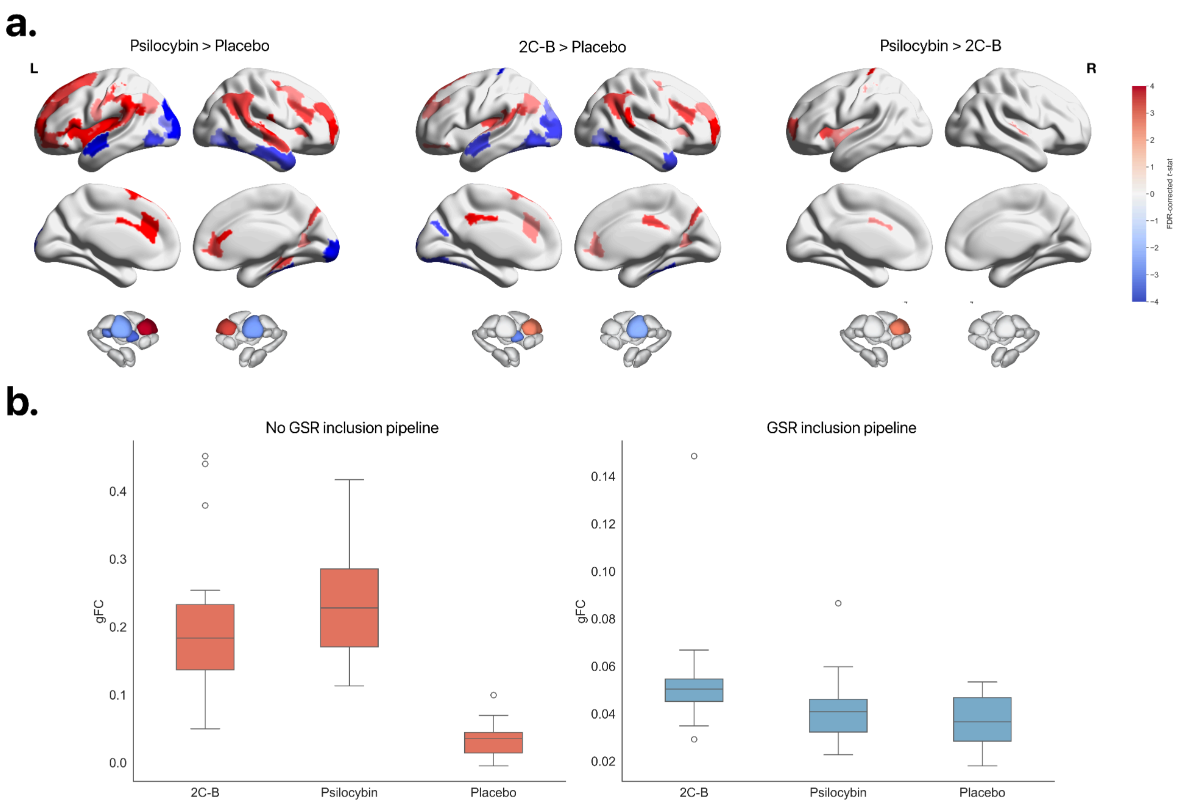


**Figure S8. Differential effects of global signal regression on global functional connectivity.** A) Significant differences in absolute GSR gFC change per condition. B) Distribution of subject whole-brain gFC values across conditions as a function of GSR inclusion.

To do so, we assessed how the absolute change in regional gFC after its inclusion may vary across comparisons. To do so, we generated absolute change maps per subject and (gFC with GSR - gFC without GSR). We performed LMEMs and corresponding follow-up tests as before, this time including framewise displacement (FD) as a nuisance covariate to control for the purported effectiveness of GSR at abating motion. As shown in Figure S8a, each contrast is affected to a different degree, with psilocybin being more affected by the use of GSR, with significant differences between 2C-B and psilocybin regarding susceptibility in prefrontal cortices and subcortical nuclei.

Similarly, we also examined how its inclusion might extend to broader whole brain gFC (Figure S8b). When assessing the interaction term of drug and pipeline choice (GSR/no GSR) while controlling for FD, we identified a significant interaction ($F_{1,115} = 62.61$) $p < 0.0001$). While all conditions were significantly influenced by its inclusion, psilocybin was disproportionately affected. Together, these findings may follow the same conclusions drawn by Avram et al, suggesting pharmacological agents are differentially susceptible to the use of GSR when excluding the influence of motion due to a range of possible latent pharmacological properties such as neurovascular coupling alterations or sympathomimetic effects (35).

### S17. References


1. Holze F, Vizeli P, Müller F, Ley L, Duerig R, Varghese N, et al. Distinct acute effects of LSD, MDMA, and D-amphetamine in healthy subjects. Neuropsychopharmacology. 2020;45(3):462-71.
2. Holze F, Becker AM, Kolaczynska KE, Duthaler U, Liechti ME. Pharmacokinetics and Pharmacodynamics of Oral Psilocybin Administration in Healthy Participants. Clinical Pharmacology & Therapeutics.n/a(n/a).
3. Dolder PC, Schmid Y, Steuer AE, Kraemer T, Rentsch KM, Hammann F, Liechti ME. Pharmacokinetics and pharmacodynamics of lysergic acid diethylamide in healthy subjects. Clinical pharmacokinetics. 2017;56:1219-30.
4. Papaseit E, Farré M, Pérez-Mañá C, Torrens M, Ventura M, Pujadas M, et al. Acute Pharmacological Effects of 2C-B in Humans: An Observational Study. Front Pharmacol. 2018;9:206.
5. Mason N, Kuypers K, Reckweg J, Müller F, Tse D, Da Rios B, et al. Spontaneous and deliberate creative cognition during and after psilocybin exposure. Translational psychiatry. 2021;11(1):209.
6. Mallaroni P, Mason NL, Reckweg JT, Paci R, Ritscher S, Toennes SW, et al. Assessment of the Acute Effects of 2C-B vs. Psilocybin on Subjective Experience, Mood, and Cognition. Clinical Pharmacology & Therapeutics. 2023;114(2):423-33.
7. Doss MK, Mallaroni P, Mason NL, Ramaekers JG. Psilocybin and 2C-B at Encoding Distort Episodic Familiarity. Biol Psychiatry Cogn Neurosci Neuroimaging. 2024.
8. Studerus E, Gamma A, Vollenweider FX. Psychometric evaluation of the altered states of consciousness rating scale (OAV). PLoS One. 2010;5(8):e12412.
9. Prugger J, Derdiyok E, Dinkelacker J, Costines C, Schmidt TT. The Altered States Database: Psychometric data from a systematic literature review. Scientific Data. 2022;9(1):720.
10. Nour MM, Evans L, Nutt D, Carhart-Harris RL. Ego-dissolution and psychedelics: validation of the ego-dissolution inventory (EDI). Frontiers in human neuroscience. 2016:269.
11. Lebedev AV, Lövdén M, Rosenthal G, Feilding A, Nutt DJ, Carhart-Harris RL. Finding the self by losing the self: Neural correlates of ego-dissolution under psilocybin. Human brain mapping. 2015;36(8):3137-53.
12. Mason NL, Kuypers KPC, Müller F, Reckweg J, Tse DHY, Toennes SW, et al. Me, myself, bye: regional alterations in glutamate and the experience of ego dissolution with psilocybin. Neuropsychopharmacology. 2020;45(12):2003-11.
13. Diaz BA, Van Der Sluis S, Moens S, Benjamins JS, Migliorati F, Stoffers D, et al. The Amsterdam Resting-State Questionnaire reveals multiple phenotypes of resting-state cognition. Front Hum Neurosci. 2013;7:446.
14. Stoffers D, Diaz BA, Chen G, den Braber A, van 't Ent D, Boomsma DI, et al. Resting-state fMRI functional connectivity is associated with sleepiness, imagery, and discontinuity of mind. PloS one. 2015;10(11):e0142014.

15. Diaz BA, Van Der Sluis S, Benjamins JS, Stoffers D, Hardstone R, Mansvelder HD, et al. The ARSQ 2.0 reveals age and personality effects on mind-wandering experiences. Frontiers in psychology. 2014;5:271.
16. Diaz BA, Hardstone R, Mansvelder HD, Van Someren EJ, Linkenkaer-Hansen K. Resting-state subjective experience and EEG biomarkers are associated with sleep-onset latency. Frontiers in psychology. 2016;7:492.
17. Mallaroni P, Mason NL, Kloft L, Reckweg JT, van Oorsouw K, Toennes SW, et al. Shared functional connectome fingerprints following ritualistic ayahuasca intake. Neuroimage. 2024;285:120480.
18. Viol A, Palhano-Fontes F, Onias H, de Araujo DB, Viswanathan GM. Shannon entropy of brain functional complex networks under the influence of the psychedelic Ayahuasca. Sci Rep. 2017;7(1):7388.
19. Schartner MM, Carhart-Harris RL, Barrett AB, Seth AK, Muthukumaraswamy SD. Increased spontaneous MEG signal diversity for psychoactive doses of ketamine, LSD and psilocybin. Scientific Reports. 2017;7(1):46421.
20. Varley TF, Carhart-Harris R, Roseman L, Menon DK, Stamatakis EA. Serotonergic psychedelics LSD & psilocybin increase the fractal dimension of cortical brain activity in spatial and temporal domains. Neuroimage. 2020;220:117049.
21. Azen R, Budescu DV. The dominance analysis approach for comparing predictors in multiple regression. Psychological Methods. 2003;8(2):129-48.
22. Wagner HH, Dray S. Generating spatially constrained null models for irregularly spaced data using Moran spectral randomization methods. Methods in Ecology and Evolution. 2015;6(10):1169-78.
23. Haenlein M, Kaplan AM. A Beginner's Guide to Partial Least Squares Analysis. Understanding Statistics. 2004;3(4):283-97.
24. Girn M, Roseman L, Bernhardt B, Smallwood J, Carhart-Harris R, Nathan Spreng R. Serotonergic psychedelic drugs LSD and psilocybin reduce the hierarchical differentiation of unimodal and transmodal cortex. Neuroimage. 2022;256:119220.
25. Luppi AI, Girn M, Rosas FE, Timmermann C, Roseman L, Erritzoe D, et al. A role for the serotonin 2A receptor in the expansion and functioning of human transmodal cortex. Brain. 2023;147(1):56-80.
26. Margulies DS, Ghosh SS, Goulas A, Falkiewicz M, Huntenburg JM, Langs G, et al. Situating the default-mode network along a principal gradient of macroscale cortical organization. Proceedings of the National Academy of Sciences. 2016;113(44):12574-9.
27. Schaefer A, Kong R, Gordon EM, Laumann TO, Zuo XN, Holmes AJ, et al. Local-Global Parcellation of the Human Cerebral Cortex from Intrinsic Functional Connectivity MRI. Cereb Cortex. 2018;28(9):3095-114.
28. Yeo BT, Krienen FM, Sepulcre J, Sabuncu MR, Lashkari D, Hollinshead M, et al. The organization of the human cerebral cortex estimated by intrinsic functional connectivity. J Neurophysiol. 2011;106(3):1125-65.
29. Alexander-Bloch AF, Shou H, Liu S, Satterthwaite TD, Glahn DC, Shinohara RT, et al. On testing for spatial correspondence between maps of human brain structure and function. Neuroimage. 2018;178:540-51.
30. Zalesky A, Fornito A, Bullmore ET. Network-based statistic: identifying differences in brain networks. Neuroimage. 2010;53(4):1197-207.
31. Murphy K, Fox MD. Towards a consensus regarding global signal regression for resting state functional connectivity MRI. NeuroImage. 2017;154:169-73.
32. Preller KH, Burt JB, Ji JL, Schleifer CH, Adkinson BD, Stämpfli P, et al. Changes in global and thalamic brain connectivity in LSD-induced altered states of consciousness are attributable to the 5-HT2A receptor. Elife. 2018;7.
33. Preller KH, Duerler P, Burt JB, Ji JL, Adkinson B, Stämpfli P, et al. Psilocybin Induces Time-Dependent Changes in Global Functional Connectivity. Biol Psychiatry. 2020;88(2):197-207.

34. Timmermann C, Roseman L, Haridas S, Rosas FE, Luan L, Kettner H, et al. Human brain effects of DMT assessed via EEG-fMRI. Proceedings of the National Academy of Sciences. 2023;120(13):e2218949120.
35. Avram M, Fortea L, Wollner L, Coenen R, Korda A, Rogg H, et al. Large-scale brain connectivity changes following the administration of lysergic acid diethylamide, d-amphetamine, and 3,4-methylenedioxyamphetamine. Molecular Psychiatry. 2024.
36. Sydnor VJ, Larsen B, Bassett DS, Alexander-Bloch A, Fair DA, Liston C, et al. Neurodevelopment of the association cortices: Patterns, mechanisms, and implications for psychopathology. Neuron. 2021;109(18):2820-46.
